# Data Matrix Normalization and Merging Strategies Minimize Batch-specific Systemic Variation in scRNA-Seq Data

**DOI:** 10.1101/2021.08.18.456898

**Authors:** Benjamin R. Babcock, Astrid Kosters, Junkai Yang, Mackenzie L. White, Eliver E. B. Ghosn

## Abstract

Single-cell RNA sequencing (scRNA-seq) can reveal accurate and sensitive RNA abundance in a single sample, but robust integration of multiple samples remains challenging. Large-scale scRNA-seq data generated by different workflows or laboratories can contain batch-specific systemic variation. Such variation challenges data integration by confounding sample-specific biology with undesirable batch-specific systemic effects. Therefore, there is a need for guidance in selecting computational and experimental approaches to minimize batch-specific impacts on data interpretation and a need to empirically evaluate the sources of systemic variation in a given dataset. To uncover the contributions of experimental variables to systemic variation, we intentionally perturb four potential sources of batch-effect in five human peripheral blood samples. We investigate sequencing replicate, sequencing depth, sample replicate, and the effects of pooling libraries for concurrent sequencing. To quantify the downstream effects of these variables on data interpretation, we introduced a new scoring metric, the Cell Misclassification Statistic (CMS), which identifies losses to cell type fidelity that occur when merging datasets of different batches. CMS reveals an undesirable overcorrection by popular batch-effect correction and data integration methods. We show that optimizing gene expression matrix normalization and merging can reduce the need for batch-effect correction and minimize the risk of overcorrecting true biological differences between samples.

## Introduction

Recent advances in throughput and commercial availability of single-cell RNA-sequencing (scRNA-seq) technology have increased accessibility and led to widespread adoption of this technology by the scientific community (Supplementary Fig. 1). A single study may encompass thousands of cells from multiple samples, often spanning time points and conditions, resulting in large and heterogenous scRNA-seq datasets^1,2^. This dramatic shift is empowering scientists to massively profile multiple samples in parallel at extremely high resolution. As experiments grow more complicated, the need arises to align and co-analyze ever larger and more diverse outputs of single-cell workflows.

Minute and often uncontrollable technical variations in sample collection and data processing can manifest as noticeable effects which confound the interpretation of data^3^. This systemic variation, referred to commonly as “batch-effect,” can pose an obstacle to data interpretation by confounding biologically-derived variation (desirable) with technically-derived variation (undesirable). An inability to discern the source of a particular signal can even lead to over-interpretation of data, in that systemic variation arising from technical differences may be interpreted as a biologically driven phenotypic difference. A typical case in which batch-effect confounds data interpretation will present as an over-merging or under-merging of cell types. Uncorrected batch-effect can cause similar cell populations between samples to appear divergent. In the inverse case, batch-effect can cause two biologically distinct populations to appear as one due to a shared technical signal. Both the prevalence and persistence of batch-specific signals have been highlighted by prior work, as well as the spectrum of methods existing to correct and remove them^4,5^. However, there remains an insufficient understanding of how experimental design and data analysis approaches play a role in producing batch-effect or identifying batch-effect when present in a sample. An understanding of the source of batch-effect and the informed selection of tools to identify batch-effect has the potential to alter the outcomes and conclusions of scRNA-seq studies.

Technical sources of variation most apparently manifest themselves in the Principal Component Analysis (PCA) matrix, representing a shared low-dimensional space. Therefore, isolation and removal of these effects in PCA dimensions are critical, as PCAs are the foundation used to produce cell cluster assignments and UMAP visualizations. Towards this aim, many methods have been applied to isolate and remove batch-effect from scRNA-seq data^6–11^. These methods try to merge biologically similar populations into a shared low-dimensional space while disregarding the influence of undesirable signals. A commonly shared assumption of current methods is that batch-specific, technically-derived signals are contained *within* the sample, while true biologically-derived signals are shared *between* samples. However, current methods are mostly agnostic to the fundamental sources of systemic variation and the underlying biological heterogeneity contained within each sample.

Here, we present a novel approach to validating batch-correction methods by demonstrating experimental variables which contribute most to systemic variation. To assess the degree of batch-effect, we introduce a biologically-grounded metric, the Cell Misclassification Statistic (CMS). While most current scoring systems are agnostic to cell identity, the CMS directly grounds itself in the cell-type classification of every single cell and is therefore uniquely able to quantify the loss of biological information during sample merging. Using CMS to quantify batch-associated systemic variation, we show that sequencing replicates and sequencing depth contribute only minimally to batch-effect and that pooling samples together for sequencing does not meaningfully improve the measured or observed batch-effect. Instead, we find that sample donor, along with the microfluidic encapsulation and library preparation steps, represent the main source of batch-associated variation. We test three popular batch-correction algorithms (Harmony^8^, LIGER^11^, and Seurat V3^10^), which have been previously scored as the best performing^5^. Our CMS scoring, which accounts for cell identity as a biological feature, revealed misclassifications of major cell lineages in all three commonly used batch-correction methods. We further applied CMS in a supervised approach to reveal that selecting a proper dataset normalization and merging strategy can perform comparably to popular batch-correction algorithms. Furthermore, we revealed that much of the batch-effect present is due to low expression levels of broadly expressed genes, which can be minimized by selecting a proper normalization and dataset merging strategy. Our unique and biologically-grounded approach allows for effective data integration in a carefully supervised workflow without the need for corrective algorithms.

## Results

### A Cell Misclassification Statistic quantifies data integration and batch-correction fidelity

We generated seven single-cell RNA-seq datasets from five individual donors of human peripheral blood mononuclear cells (PBMCs) (Fig. 1a). To quantify the contributions of known variables to batch effect and evaluate the performance of data integration and batch correction methods, we developed a cell misclassification statistic (CMS) metric (Fig. 1b). CMS is grounded in the premise that if different systems identify a single cell as different biological types, both cannot be correct. To calculate a CMS, we first gather the cell-type identities obtained by classifying cells from only a single sample, meaning that all cells were processed under identical conditions and are influenced identically by potential technical sources of variance, if at all. Then, to calculate a CMS score, we compare the cell-type classifications of each sample individually to cell-type classifications after multiple sample merging. We measure the fraction of cells that have changed classification and generate a statistic, such that a CMS score of 0 means that no cells changed classification/cell-type identity after merging datasets, while a CMS of 0.2 means a misclassification of 20% of cells. A higher CMS score will result when cell barcodes change cell-type identity after sample merging and indicates over-correction of the sample. By relying on invariable biological principles (a single, non-doublet cell barcode must hold only one cell classification), we directly interpret CMS scores as a measure of biological signal loss during data integration.

**Fig. 1:**
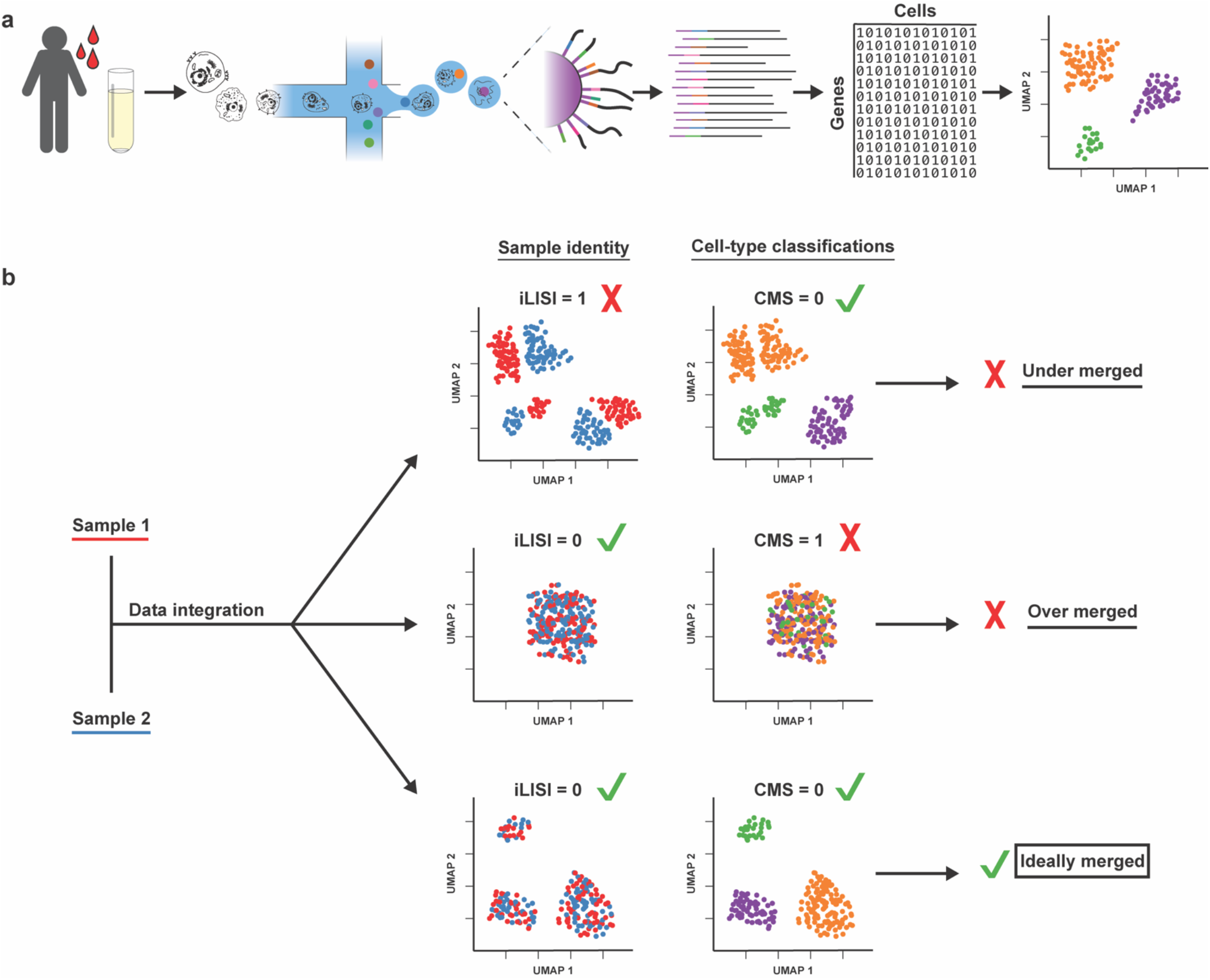
Schematic of analysis workflow to quantify data integration and batch-correction fidelity. **a,** Tissue (blood) is extracted from the sample donor, washed, and PBMCs isolated. PBMCs are encapsulated in microfluidic droplets, along with a barcode-bearing bead. Cells are lysed in the droplets, and mRNA is captured on the bead, resulting in a barcoded cDNA library. Libraries are sequenced to generate a GEX matrix (cells x genes) containing transcript counts. Cells are analyzed and plotted by UMAP and clustered according to transcript similarity, after which cell types are classified. **b,** Cell Misclassification Statistic (CMS) and modified integration LISI (iLISI) metrics provide measures of cell-type fidelity and UMAP mixing, respectively. CMS and iLISI can assist in optimizing a workflow for the best-mixed UMAP while preserving the cell-type-specific signals in the data.

To produce the initial cell type classifications required for CMS, we analyzed seven datasets generated from five PBMC samples. Each dataset contained two matrices holding independent modalities of data: gene expression (GEX) and antibody-derived tag (ADT) cell-surface protein expression. First, we confirmed the cell types present in each single sample by removing an aliquot of cells and performing flow cytometry staining and analysis by a gold-standard “gating strategy” approach (representative data shown in Supplementary Fig. 2). In parallel, we processed the GEX sequencing data to generate clusters according to the Seurat V3 workflow, following default settings^10^. We next assigned each cluster to a major lineage using both ADT and GEX markers based on a gating strategy similar to flow cytometry, as we have previously reported^12^ (representative data shown in Supplementary Fig. 3). With cell-type classifications independently established for each sample, we can assess the impacts of batch-effect on cell-type classification in merged datasets.

CMS measures consistency in cell-type classification, which is only one of two metrics we used to validate data integration. Another important aspect of data integration and batch correction methods is generating UMAP embeddings for data visualization. To assess the performance of UMAP embeddings for visualization of integrated datasets, we employed a modified Local Inverse Simpson’s Index (LISI)^8,13^, which we used specifically to measure the final UMAP integration, or integration LISI (iLISI) (Fig. 1b). In our system, an iLISI score of 1 represents a UMAP completely segregated by sample ID, while an iLISI of 0 represents a perfectly integrated UMAP. Therefore, the best-integrated data will have both CMS and iLISI scores approaching 0, while the most segregated data will return both CMS and iLISI scores approaching 1 (Fig. 1b). As described below, we applied the CMS and iLISI methods to compare the fidelity of the various data-integration/batch-correction methods we evaluated in this study.

### Batch-associated systemic variation is observed when sample replicates and sequencing depth are perturbed, but not sequencing replicate

To examine the contributions of experimental variables to batch-associated data effects, we first describe four unique steps of a typical scRNA-seq workflow in which sample-specific variation may be introduced:

1. Sample donor
2. Sample and library preparation
3. Library sequencing
4. Data analysis

The sample donor (i) represents the baseline variation intrinsic to each human subject who donated blood for this study. Sample preparation (ii) includes the processes of extracting cells from donor tissue and preparing a single-cell suspension (the sample) (Fig. 1a). Together, sample and library preparation (ii) encompasses all steps spanning microfluidic encapsulation of donor cells through to the generation of a barcoded cDNA library (Fig. 1a). (Note: As sample and library preparation will largely be tissue and platform-specific and will not typically be varied for a single-donor aliquot of cells, we consider them here as processes intrinsically linked to sample donor and not as independent sources of batch-effect). Library sequencing (iii) consists of converting the cDNA molecules (i.e., barcoded libraries) into aligned reads, and finally, an expression matrix of gene counts per cell (Fig. 1a). Data analysis (iv) constitutes the last unit of the workflow. During data analysis, we make interpretations on the expression matrix by assigning cell type IDs to cell barcodes and clusters, visualizing cells by UMAP embeddings, and detecting differential gene expression (DGE) (Fig. 1a). To evaluate the contributions of each of these steps (i-iv) to batch-effect, we knowingly introduced differences in key variables to sets of human PBMC samples and evaluated the downstream batch-effect observed. Finally, we directly evaluated whether pooling libraries for concurrent sequencing could reduce sample-associated batch-effect compared to the same libraries sequenced independently.

#### Sample donor

We assessed the systemic batch variation in identically-prepared PBMC samples from three healthy adult donors (samples 1-3, Supplementary Table 1). We processed samples simultaneously for PBMC isolation, single-cell encapsulation, and barcoded cDNA library generation. Following library preparation, we pooled samples for simultaneous sequencing. Depth of sequence (reads per cell) was not significantly different between samples (Wilcoxon rank-sum test, p = 0.3487, 0.2445, 0.8471 for samples 1, 2, 3, respectively), and we proceeded with 16,946 cell barcodes after concatenating the GEX matrices. Next, we assigned each cluster to a cell type using both GEX and ADT data (Supplementary Fig. 3). After sample merging, the three PBMC samples were mostly segregated by donor when visualized on a UMAP (Fig. 2a). The failure to effectively integrate the three samples, as readily apparent in the UMAP embedding, was confirmed by our modified iLISI scoring (iLISI = 0.861, Fig. 2a). We then applied CMS scoring and revealed approximately ten percent cell-type misclassification after sample merging (CMS = 0.099, Figs. 2a and Supplementary Fig. 4). Hence, we demonstrate that sample-specific variations cause a loss of biological signal in the integrated data analysis and contribute to undesirable batch-associated data effects.

**Fig. 2:**
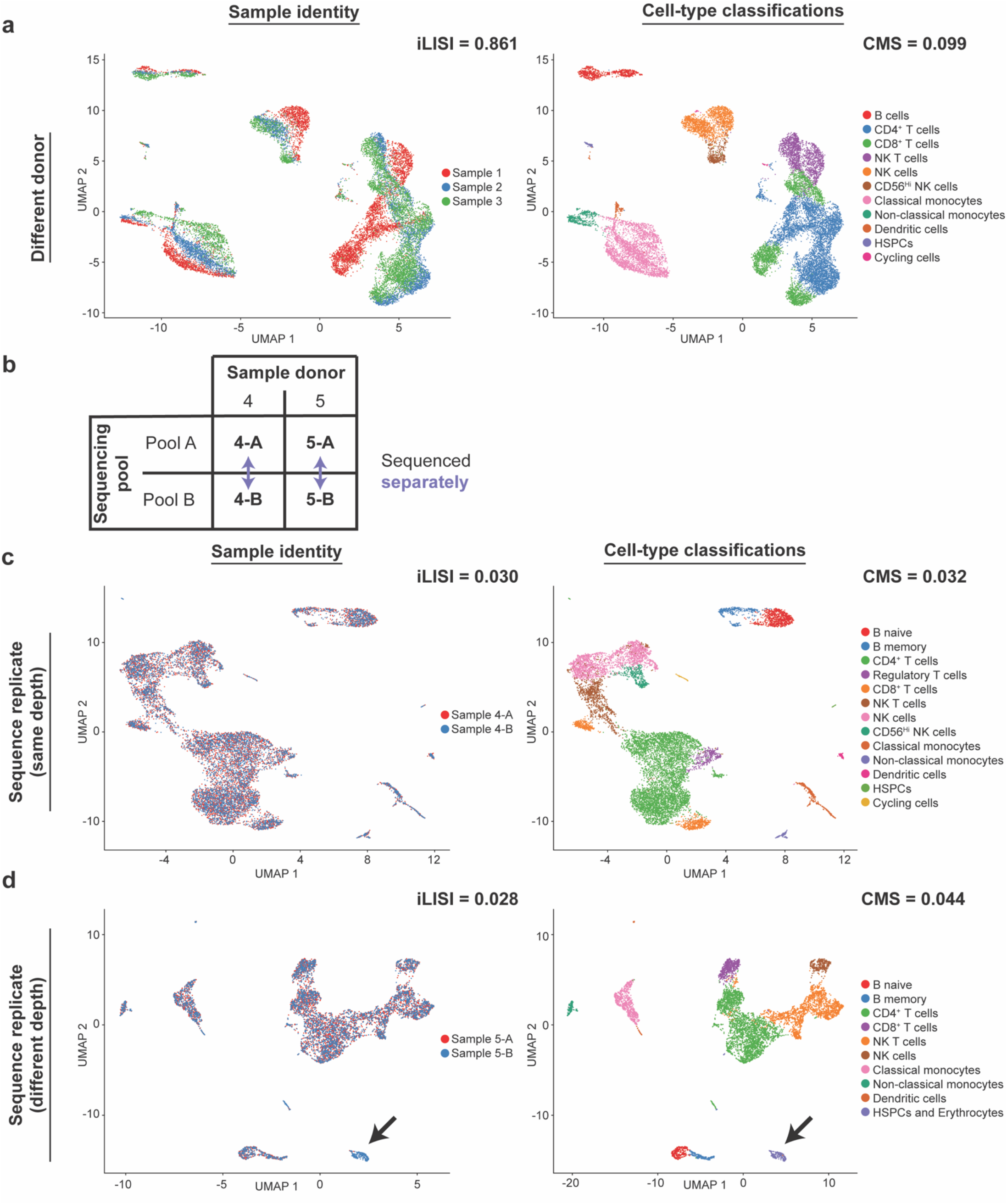
Batch effects are generated by sample donor and sequencing depth, but not sequencing replicates alone. **a,** PBMCs from three different donors, processed and sequenced simultaneously and mixed for co-analysis, produce a favorable cell-type classification fidelity (CMS = 0.099) but an unfavorable (i.e., poorly mixed) UMAP (iLISI = 0.861). **b,** A schematic of sample sequencing strategy. Libraries from two donors (samples 4 and 5) were sequenced twice. **c,** A library generated from a single PBMC sample, divided and sequenced twice to similar read depth (average reads per cell), produces a well-integrated UMAP and high-fidelity cell classifications (iLISI = 0.030, CMS = 0.032). **d,** A library generated from a single PBMC sample, divided and sequenced twice with a three-fold difference in read depth produces favorable UMAP and cell-type fidelity metrics (iLISI = 0.028, CMS = 0.044). However, there are local sample-specific UMAP islands (black arrow) and misclassified cell types that reveal sequencing-depth-specific batch effects.

#### Library Sequencing Replication

To demonstrate the contributions of library sequencing to systemic variation, we aliquoted a single library in two and performed duplicate sequencing under similar conditions (samples 4-A and 4-B) (Fig. 2b). We sequenced each aliquot in a separate flow cell, on different days, with similar target sequence depth (∼40,000 reads/cell) (Fig. 2b). After merging, the dataset contained approximately 13,000 cells, representing a shared set of approximately 6,500 barcodes, duplicated. We found that 6,392, or greater than 99 percent, of cell barcodes, were shared between both sequencing replicates, indicating minimal new cells were identified by additional sequencing. As above, we followed a default Seurat workflow to generate UMAP visualizations and Louvain clusters (Fig. 2c) and used the GEX and ADT data to classify clusters by cell type. We constructed a table of the proportion of each sample that contributed to each cell type and cluster (Supplementary Table 2). As both libraries are derived from two aliquots of the same cDNA pool, we would expect the proportions of cell types to be equal by replicate, which they were (Supplementary Table 2; chi-square test, p = 1). Scoring the replicates by iLISI confirmed a homogenous distribution of replicates in the UMAP (iLISI = 0.030) while CMS revealed only 3.2% of cells changed classification (CMS = 0.032, Supplementary Fig. 4), which represents fewer cells than the expected cell-cell doublet rate for this technology. Hence, we find that variation introduced by library sequencing was a non-significant contributor to batch-effect by all measures.

#### Library Sequencing Depth

Next, we quantified the effect of sequencing depth by intentionally altering the number of reads between replicates. For this comparison, we constructed two identical aliquots of a PBMC library containing approximately 4,000 cells, sequenced to different depths (samples 5-A and 5-B) (Fig. 2b). Sample 5-B contained three-fold more total reads than sample 5-A (170M reads for 5-A vs. 560M reads for 5-B). Notably, we sequenced samples 5-A and 5-B on the same Illumina flow cell as samples 4-A and 4-B, respectively, from the experiment above (Fig. 2b). Having already established that library sequencing replication (i.e., sample 4) produced minimal batch-effect (Fig. 2c), we interpret that any differences observed between samples 5-A and 5-B can be attributed to sequence depth. We recovered similar numbers of cells per replicate, resulting in averages of 45,160 reads/cell in sample 5-A and 138,274 reads/cell in 5-B, with 94% overlap in cell barcode sequences (3,758 of 4,007). Of the 249 non-shared cell barcodes, all were present only in 5-B, the sample with greater sequencing depth (i.e., higher reads/cell). As before, we analyzed the gene expression matrices following the Seurat workflow under default settings and assigned cell types using a GEX and ADT-based gating approach (Fig. 2d). Surprisingly, in this comparison cell-type compositions were biased by sequencing replicate (Supplementary Table 2; chi-square, p < 2.2×10^−16^) while iLISI and CMS scores remained favorable (iLISI = 0.028, CMS = 0.033). One notable cell type, erythrocytes, stood out as batch-biased: 0.3% of total cells in sample 5-A were classified as erythrocytes, compared to 5% in 5-B. Indeed, removing the erythrocyte-classified population resulted in a homogenous composition of cell types by sample (Supplementary Table 2; chi-square, p = 1). Removing erythrocytes also improved the CMS score from 0.033 to 0.017, still well below the expected cell-cell doublet rate.

We hypothesized that erythrocyte-classified barcodes appeared only in the greater read depth sample because they contain fewer unique molecular identifiers (UMIs), requiring more reads to capture their relatively rare transcripts. We confirmed that erythrocytes did contain fewer unique mRNA molecules (Supplementary Fig. 5, Wilcoxon rank-sum test, p < 2.2×10^−16^). In addition, we observed that hematopoietic stem and progenitor cells (HSPCs) co-clustered with erythrocytes, which may explain the few high-UMI dots present in Supplementary Fig. 5. We also confirmed that sample 5-B (the higher read-depth sample) contained more UMIs per cell than sample 5-A (Supplementary Fig. 5, Wilcoxon rank-sum test, p = 1.482×10^−10^), and that sample 5-B also contained more unique genes per cell (Supplementary Fig. 5, Wilcoxon rank-sum test, p < 2.2×10^−16^). As a standard part of the 10X Genomics Cell Ranger workflow (i.e., the sequencing read alignment and demultiplexing steps), cell barcodes with low-UMI counts are excluded as potential empty droplets containing ambient mRNA/noise. To confirm that the erythrocyte cell barcodes unique to sample 5-B were also present in the aliquot sequenced for 5-A but instead had been artificially excluded as low-UMI barcodes, we investigated the unfiltered sequencing matrices of sample 5-A. We discovered 100% of cell barcodes specific to 5-B were contained within the unfiltered 5-A data matrix but were excluded as part of the Cell Ranger quality control steps. Only in sample 5-B, the high-depth replicate, were enough UMIs sequenced to distinguish erythrocytes from ambient noise. Hence, our results demonstrate that imbalanced sequencing depth may result in cell type biases, contributing somewhat to batch-associated data effects.

#### Pooled Library Sequencing

It is widely assumed that technical effects can be minimized by pooling libraries for sequencing in a single batch. However, we demonstrate above that duplicated sequencing of identical library aliquots can yield highly similar results with minimal batch-specific variation (Fig. 2c). To directly evaluate the benefits of sequencing libraries together in a single pool, compared to sequencing un-pooled libraries independently, we again analyzed the sequenced replicates of PBMC samples 4 and 5 (samples 4-A, 4-B, and 5-A). We do not expect samples 4 and 5 to be identical, as they originate from different sample donors. However, we will directly contrast the analysis of merged samples 4-A/5-A, which we sequenced in a single pool, against the analysis of merged samples 4-B/5-A, which we sequenced in different pools (see experimental design, Fig. 3a). If indeed pooled sequencing alleviates batch-effect, we would expect to identify less batch-specific variation (lower iLISI and CMS scores) when merging samples that were pooled for sequencing together (samples 4-A vs. 5-A) as compared to un-pooled samples sequenced separately (samples 4-B vs. 5-A). Notably, we sequenced all samples for these comparisons to a highly similar read depth (∼40,000 reads/cell). Surprisingly, we find that pooling libraries for sequencing yields nearly identical results as sequencing unpooled libraries independently (Figs. 3b-c). Cell misclassification rates rose compared to the previously presented experiments of duplicated sequencing of identical libraries (Fig. 2d) but remained constant between pooled and un-pooled comparisons (CMS = 0.070 and 0.070, respectively). The UMAP homogeneity similarly remained near-constant between pooled and un-pooled sequencing batches (iLISI = 0.279, 0.295, respectively; Figs. 3b-c). Our results assert that pooling libraries for sequencing neither reduced nor contributed to a major source of batch-associated data effects in these samples.

**Fig. 3:**
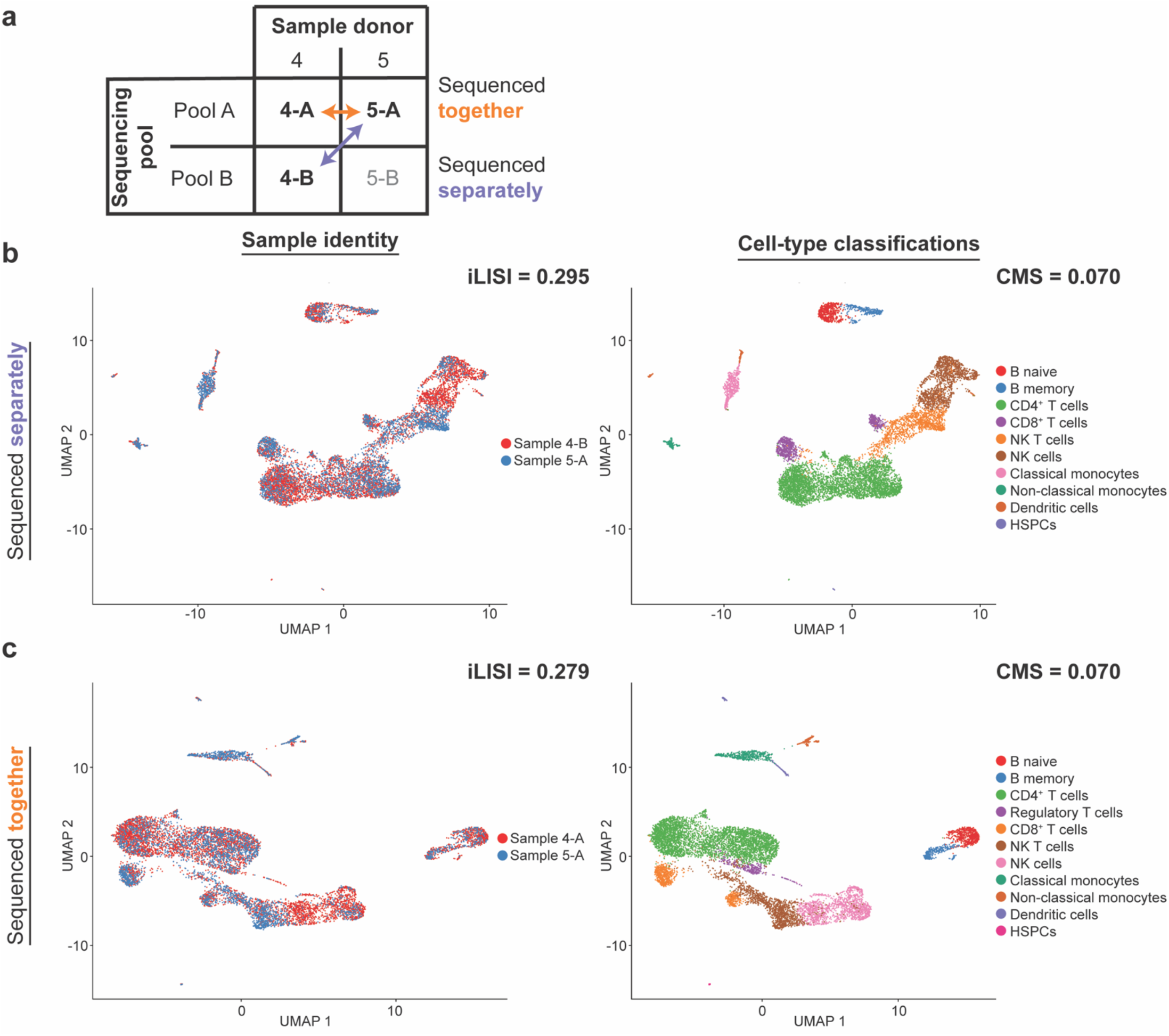
Pooling samples for sequencing does not appreciably improve the measured batch effect. **a,** A schematic of pooling and comparison strategy between samples 4 and 5, sequenced across two independent pools (-A and -B). The same sample (5-A) was co-analyzed with either a sample from the same sequencing pool (4-A) or a different pool (4-B). **b,** Analysis of two PBMC samples from different sequencing pools results in a poorly mixed UMAP and the misclassification of 7% of cells (iLISI = 0.295, CMS = 0.070). **c,** Pooling libraries to sequence PBMC samples in a single pool, followed by mixing data for co-analysis, does not improve batch effects and, similarly, results in a poorly mixed UMAP and an equivalent 7% cell misclassification rate (iLISI = 0.279, CMS = 0.070).

### Commonly used batch-effect correction and data integration methods imperfectly resolve batch-associated systemic variation

We examined the extent to which commonly cited data integration algorithms can minimize the batch-specific systemic variation we quantified in the prior experiments. Above, we showed that the sample donor variable significantly confounded our ability to correctly assign cell types and resulted in a sample-biased UMAP from a merged dataset (samples 1-3; Fig. 2a, iLISI = 0.861). Here, we attempt to improve the low-dimensional embeddings (i.e., UMAP and clusters) and the robustness of cell-type classifications by applying commonly-used batch-effect correction algorithms. First, we selected three of the most cited packages for integrating gene expression data: Harmony^8^, LIGER^11^, and Seurat V3 (SCTransform and data anchoring)^6,10^, which have been established as the best performing by prior study^5^. Next, we followed the default workflow settings recommended by the respective publications and applied each method to our PBMC samples 1-3. Finally, we generated UMAP embeddings and cluster/cell-type assignments to assess each corrected dataset (Fig. 4).

**Fig. 4:**
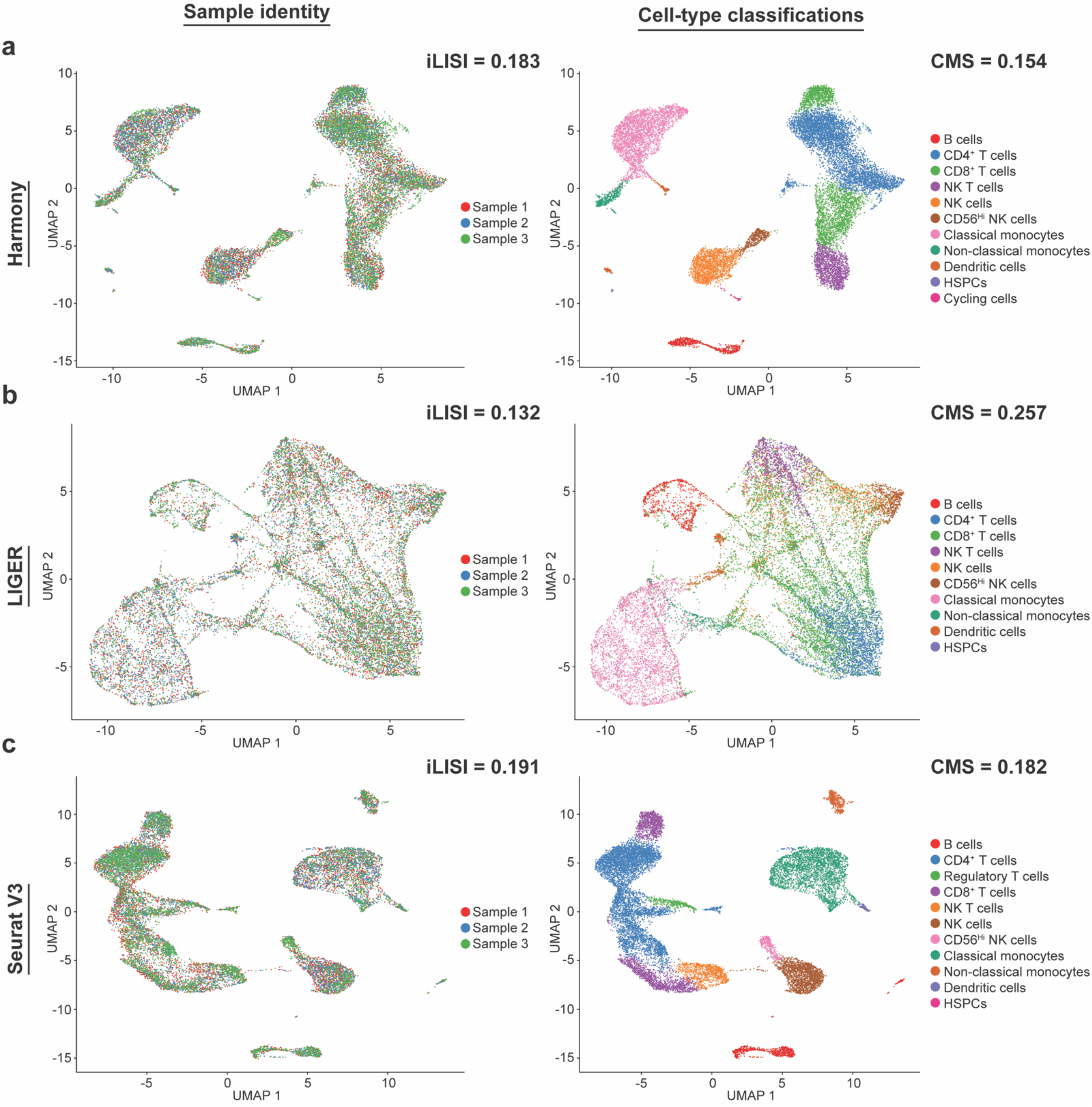
Common batch-effect correction methods imperfectly resolve batch effect. **a,** Harmony correction merges samples to produce an integrated UMAP, but at 15.4% loss in cell-type classification fidelity (iLISI = 0.183, CMS = 0.154). **b,** LIGER correction produces an integrated UMAP, but at a 25.7% loss in cell-type classification fidelity (iLISI = 0.132, CMS = 0.257). **c,** Seurat V3 correction results in an integrated UMAP but at a cost of an 18.2% loss in cell-type classification fidelity (iLISI = 0.191, CMS = 0.182).

We first evaluated Harmony, implemented through the Seurat function “RunHarmony”^8,14^. Although homogeneity of the UMAP is greatly improved (Fig. 4a; iLISI improved from 0.861 to 0.183), CMS scoring of the harmonized sample revealed a greater loss of cell-type fidelity (Fig. 4a; CMS rose from 0.099 to 0.154). Interestingly, the increased proportion of cell-type misclassification (from 9.9% in the uncorrected sample to 15.4% in the harmonized sample) comes largely from over-merging CD4^+^ and CD8^+^ T cells into mixed-lineage clusters (Figs. 4a and Supplementary Fig. 4). This highlights the potential to over-homogenize data while ignoring specific cell type-exclusive signals.

We next tested LIGER, as implemented through the R package “rliger” and the Seurat functions “RunOptimizeALS” and “RunQuantileNorm”^11,15,16^. We generated a UMAP from the LIGER integrative non-negative matrix factorization (iNMF) components (Fig. 4b). LIGER produced a homogeneously distributed UMAP (Fig. 4b; iLISI = 0.132) but severely over-merged clusters, resulting in a major loss of cell-type information for nearly one-quarter of all cells, as measured by a severely increased CMS score (Fig. 4b; CMS rose from 0.099 to 0.257). As with Harmony, the most affected cell types were the CD4^+^ and CD8^+^ T cells. LIGER could not differentiate those two distinct subsets of T cells (Supplementary Fig. 4) and incurred a misclassification cost of greater than 25% of cells (i.e., CMS = 0.257).

Lastly, we evaluated the performance of Seurat V3, as implemented through the Seurat SCTransform and data anchoring workflows^6,17^. Data integration by Seurat V3 resulted in improved homogeneity of UMAP embeddings (Fig. 4c; iLISI improved from 0.861 to 0.191), accompanied by an increase in CMS scores compared to uncorrected data (Fig. 4c; CMS rose from 0.099 to 0.182). Again, as with both Harmony and Liger, the Seurat data integration methods were unable to effectively segregate subsets of CD4^+^ and CD8^+^ T cells (Supplementary Fig. 4). We conclude that similar yet transcriptionally distinct types of PBMCs pose a problem for data integration methods as all three failed to resolve these specific T-cell subsets. While we focus our subsequent efforts on detailing the batch-effect impacting T-cell subsets, it is important to note that these effects were global and impacted other major lineages, including NK cells, B cells, and monocytes (Supplementary Fig. 4). Crucially, each method generated a different set of clusters when applied to the same merged dataset (Fig. 4), highlighting a potential to misinterpret results from a single method, especially when relying on clusters as a meaningful descriptor of biological status. Notably, by applying the CMS score, we are uniquely able to reveal and quantify the hazards that dataset integration can pose to data interpretation.

### Optimized dataset normalization and scaling before integration resolves batch-effect without the need for batch-correction methods

We showed that batch-specific variations negatively impact UMAP embeddings (iLISI scores) and cell-type classifications (CMS scores) in merged samples, and yet commonly-used batch-correction methods imperfectly resolve these effects (Figs. 2 and 4). Hence, we sought to identify and optimize the specific data processing steps which can influence downstream UMAP embeddings and clustering. We focused on the steps of data normalization and data scaling, which are required to produce comparable transcript counts between cells and across genes, and must be applied prior to any PCA and downstream UMAP and clustering. To quantify the effect of data normalization and scaling on resolving batch-associated systemic variation, we applied two unique data normalization and scaling methods with two different data pooling workflows to generate a total of four datasets for comparison (Fig. 5).

**Fig. 5:**
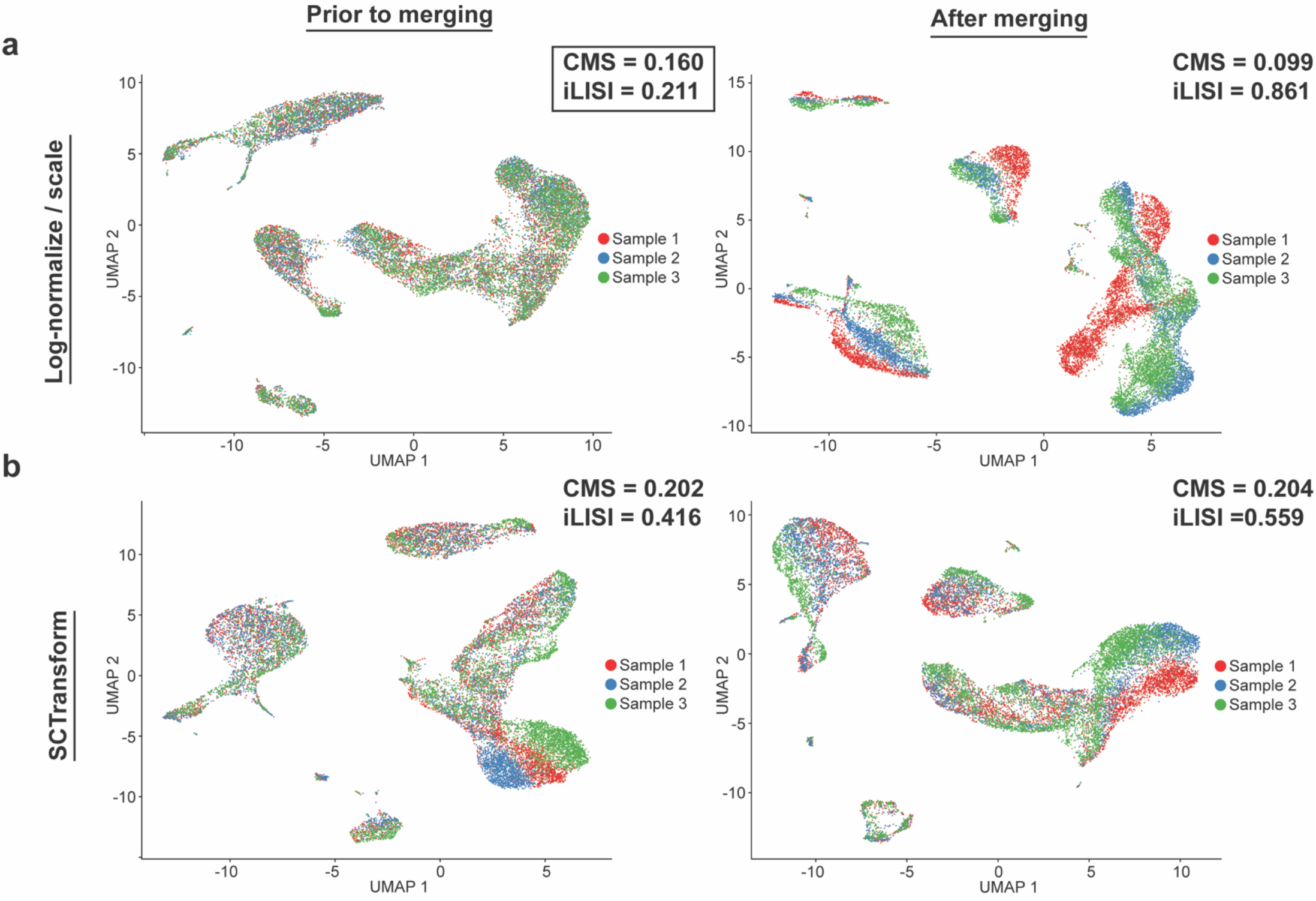
Data normalization and merging strategies differentially impact the measured batch effect. **a,** UMAPs, colored by sample, when the same normalization methods (log-normalization and data scaling) are performed on each sample either prior to sample merging (i.e., normalized separately) or after sample merging (normalized together). **b,** UMAPs, colored by sample, when the SCTransform normalization method is performed before or after sample merging. For this data set, log-normalization/scaling performed on each sample separately, prior to sample merging, generated the best combination of both integration scores, CMS and iLISI (panel a, left).

We first evaluated the common normalization/scaling method of log-transformation and gene centering, as implemented by Seurat methods NormalizeData and ScaleData^10^. Because each cell has a different number of UMI sequenced, NormalizeData divides gene count values by the total number of read counts per cell and multiplies by a scaling factor (10,000 by default). The result is a scaling of each cell to a total of 10k UMIs to avoid the effect of a different sequencing depth across cell types in a sample. NormalizeData then adds a pseudocount of 1 (to avoid transcript zero-values) and takes the natural log of each count. In this way, we normalize cells for per-cell sequencing depth, fostering more similar cell-cell comparisons. The data requires further scaling, however, to stabilize the relationship between gene expression level and variance. ScaleData employs a simple gene-level centering and scaling, meaning that each gene will be mean-centered to zero and expression values scaled by the standard deviation. The resulting scaled values (a z-score) are clipped to a maximum (default of 10) to reduce the effect of outlier high-variance genes expressed by a minority subset of cells.

The second method evaluated is the SCTransform method included in Seurat V3^10^. Briefly, SCTransform models UMI counts using a regularized negative binomial model to remove the cell-cell variation caused by differing sequencing depth between cell barcodes. In accomplishing this, SCTransform pools genes with similar abundance to obtain stable parameter estimates, preventing the overfitting caused by a global scaling model. In this way, SCTransform simultaneously corrects for influences of both total UMI and mean expression on the gene variance.

We applied the above two normalization/scaling methods to PBMC samples 1-3, which had the greatest batch-effect in our prior experiments (Fig. 2a), and which were ineffectively integrated by common batch-effect correction algorithms (Fig. 4). Importantly, each dataset was processed by each method, either log-normalization and scaling (Fig. 5a) or SCTransform (Fig. 5b), both prior to data merging or after data merging as a single unified/concatenated dataset. We then assessed the differences in the final UMAP visualization and cluster compositions and generated iLISI and CMS scores for each method (Figs. 5 and Supplementary Fig. 4). SCTransform normalization produced undesirable results, showing greater numbers of cell-type misclassification (high CMS scores) and sample-stratified UMAP embeddings (high iLISI scores), irrespective of whether it was performed prior to or after dataset merging (Fig. 5b; CMS = 0.202 and iLISI = 0.416 prior to merging; CMS = 0.204 and iLISI = 0.559 after merging). We identified one method, log-normalization with mean-centering performed on each sample independently and prior to data merging, that produced the best homogenized UMAP (Fig. 5a; iLISI = 0.211). However, nearly one in every six cells was misclassified after dataset merging (Fig. 5a; CMS = 0.159). In contrast, the same normalization/scaling method performed after dataset merging showed a highly stratified UMAP embedding, even though it had the best CMS score (Fig. 5a; iLISI = 0.861, CMS = 0.099). The simple default methods of log-normalization with mean-centering performed prior to dataset merging surprised us by their ability to nearly match the performance of dedicated batch-effect correction algorithms (Figs. 4 and 5a).

Taken together, our results implicate data normalization and scaling as an effective method to de-emphasize the systemic variation present in the sample-specific data matrix, performing similarly to the Harmony and Seurat V3 batch-effect correction methods. However, even the best normalization approaches led to high levels of cell misclassification, which carry a dangerous potential for misinterpretation of data and false conclusions.

### Low levels of highly variable transcripts are associated with batch-effect

We show above that normalization and merging strategies can have dramatic impacts on the observed levels of batch-effect. Here, we attempt to isolate the gene transcripts, which may be responsible for the batch effects observed when the integrated samples are normalized differently.

Zero-count gene transcripts can indicate either the true absence of a transcript, or that the expression level of a given transcript is very low and may not be captured by the assay, resulting in what is known as gene “dropout” events. Both true zero-count transcripts and gene dropouts account for a large proportion of the gene expression matrix and can contribute significantly to batch-effect^18,19^. One key difference between general normalization methods commonly used for bulk RNA-seq and those specifically developed for scRNA-seq is the ability to cope with excessive zeros in the scRNA-seq data matrix^18^. Hence, we reasoned that low expression level transcripts (fewer than 5 UMI) may be susceptible to gene dropout (i.e., zero-count transcripts) and that proper normalization may minimize the effect of these low expression level genes.

To identify precisely how normalization methods could improve cell-type classification, we first isolated the misclassified cells of sample 1, the individual sample with the greatest total cell-type misclassification (i.e., highest CMS score) of the PBMCS samples 1-3 (Figs. 2a, 4, and 5). We found that a specific group of cells in sample 1, the CD8^+^ T cells, frequently changed their initial classification when the cell-typing workflow to classify clusters (Supplementary Fig. 3) was repeated/validated after sample merging. This group of CD8+ T cells was prone to inappropriately co-cluster with NK cells, along with another group of CD8+ T cells which were co-clustered with CD4^+^ T cells and regulatory T cells (Fig. 6a). Notably, this observation was not exclusive to a single computational workflow (Supplementary Fig. 4) and was repeated across all normalization methods tested (Fig. 5).

**Fig. 6:**
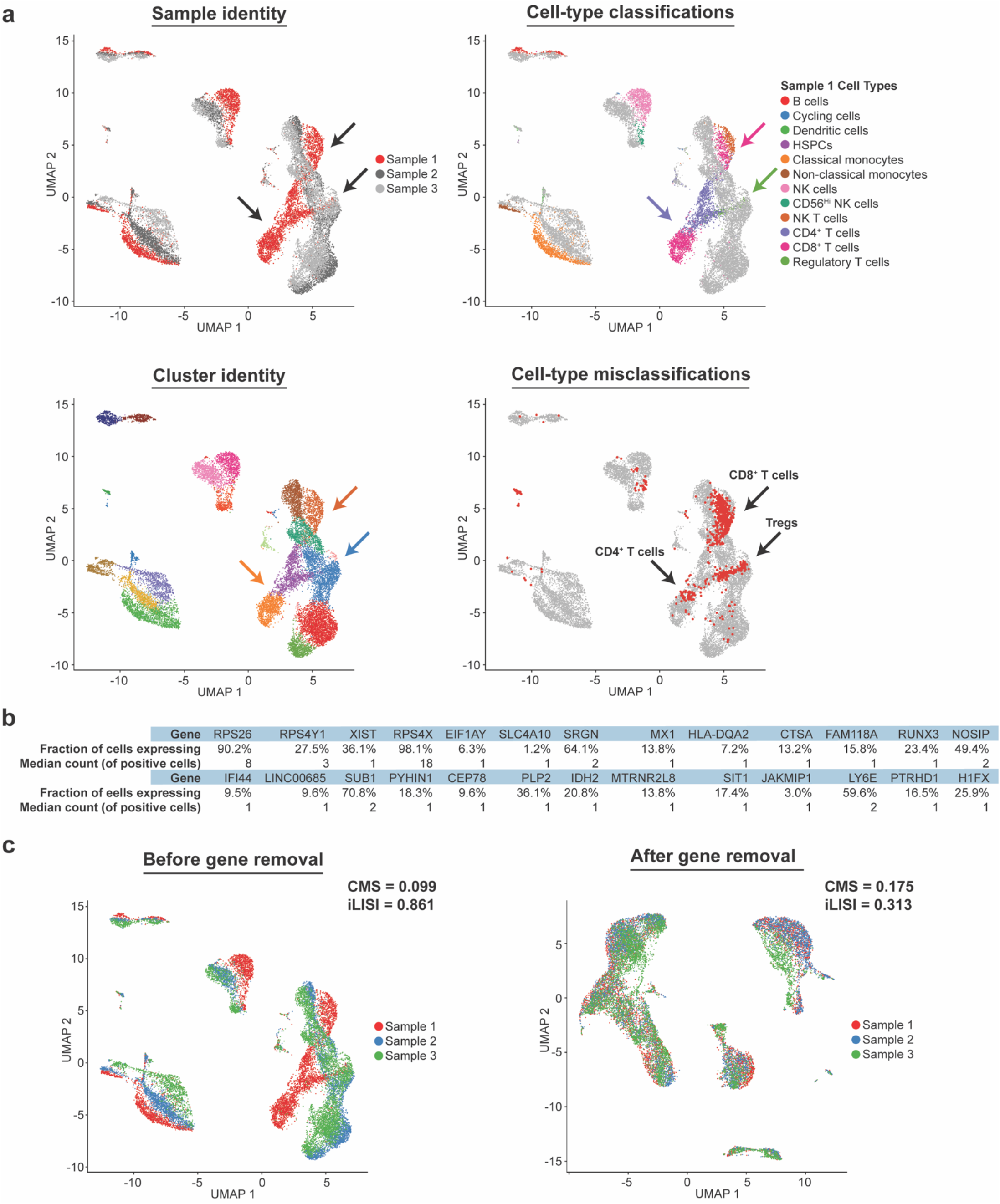
Low expression levels of highly-variable genes can play a role in batch effect, which can be amplified by the data normalization and merging strategies. **a,** Misclassified cells are not distributed at random, and specific analysis of PBMC sample 1 reveals a loss of cell-type identity resulting from the over-merging of T cell subsets into clusters containing a majority of a different subset (misclassified T cells highlighted by arrows and red points). **b,** Selected genes identified as differentially expressed between the incorrectly classified CD8^+^ T-cells of sample 1, marked by an arrow in (a), compared to the correctly classified CD8+ T-cell cluster. Genes were restricted to those contained within the HVGs of sample 1, but not the HVGs of samples 2 or 3. Note that genes showing low expression levels (low median count) and broad expression (higher fraction of cells) are susceptible to gene dropout. **c,** Removal of selected genes (b) lessens but does not completely remove the batch effect from the sample (iLISI changes from 0.861 to 0.313, CMS changes from 0.099 to 0.175).

We next sought to identify the specific gene transcripts which can distinguish the misclustered CD8+ T cells of sample 1 from the CD8+ T cells of samples 2 and 3. We directly performed differential gene expression (DGE) analyses between the misclassified CD8+ T cells of sample 1 vs. the correctly classified CD8+ T cell cluster using a likelihood ratio test for differential expression, as implemented by Seurat “FindMarkers” function^10,20^, and identified a set of significant differentially-expressed genes (Fig. 6b). Next, to restrict this list to genes that could directly influence the UMAP and clusters, we retained only genes detected as “highly variable genes” (HVGs). Only HVGs are included as input to the PCA, which is used as the input for the downstream UMAP and clustering. Finally, to assess whether the genes in our list could cause batch effect, we excluded the genes that were in the HVGs of the dataset that showed the least batch effect (Fig. 5a, left panel). The final list is a limited set of genes that are differentially expressed between the CD8^+^ T cells of sample 1 and the samples 2/3, are capable of influencing PCA (and downstream UMAP and clustering), and are present in the HVGs of datasets which show strong batch effect. This final list of 26 genes is presented in Fig. 6b.

For 23 out of 26 genes present in this final list, the median transcript count (of cells with nonzero transcripts) was two or fewer copies (i.e., ≤ 2 UMI). The more highly expressed genes were all ribosomal subunits (*RPS26*, *RPS4Y1*, *RPS4X*), which had median transcript counts of 8, 3, and 18 UMIs, respectively (Fig. 6b). Remarkably, we show that specific removal of these genes from the HVGs of samples 1-3 resulted in a marked improvement in data integration without the need for any other optimization (Fig. 6c). iLISI scores improved from 0.861 to 0.313, and CMS scores approached the scores achieved by the best batch-correction and normalization approaches (CMS = 0.175). Therefore, our results show that the selection of normalization/scaling and sample merging workflow plays an important role to either exacerbate or minimize batch-associated systemic variation by properly controlling contributions of low-expression genes to the total variance of the sample. In conclusion, here, we experimentally demonstrated the various sources of batch-associated effects and developed a new scoring system that takes into account cell-type identity as a key biological feature in data integration efficiency. Taken together, these results can inform the optimal experimental design, and data integration approaches.

## Discussion

As accessibility to scRNA-seq dramatically increases, so too do the challenges of integrating and co-analyzing diverse datasets. One challenge to integrating multiple scRNA-seq samples is the influence of batch-specific technical variance on UMAP visualization and clustering. Here, we apply our novel Cell Misclassification Statistic (CMS) score alongside a modified Local Inverse Simpson’s Index (iLISI) scoring to reveal previously unclarified sources of these batch-specific effects and the potential dangers of using batch-correction algorithms. We conclude that batch-effect can be partially mitigated by supervised optimization of normalization and scaling methods, which work by minimizing the influence of low-expression gene transcripts.

Existing batch-correction scoring metrics such as Local Inverse Simpson’s Index (LISI)^8,13^ or Average Silhouette Width (ASW)^21^ quantify the mixing of merged samples in either the integrated clusters or in the final integrated UMAP. However, these and other current batch-correction approaches remain agnostic to the biology of each cell, such that potential cell-type misclassification after data integration is not considered in the analysis. Yet, as we show here, cell-type misclassification is a pervasive phenomenon in data integration and can dramatically influence data interpretation. Therefore, we contend that in addition to measuring optimal mixing of integrated samples (e.g., LISI or ASW), the potential for cell-type misclassification post sample merging must be taken into account when evaluating the fidelity of batch-correction and data integration methods.

In this study, we developed the CMS score to directly measure cell-type misclassification that often occurs following batch correction and data integration. CMS differs from existing approaches such as LISI^8,13^ or ASW^21^ in that CMS quantifies batch-integration based on a known biological classification of cells, rather than by simply measuring the mixing of different samples. CMS is uniquely sensitive to the incorrect merging of dissimilar cell types caused by the over-homogenization of datasets. To complement these strengths, we combine CMS with a modified LISI score, or the integration LISI (iLISI), to measure the efficiency of the UMAP integration. iLISI is uniquely sensitive to detecting the under-merging of similar samples, a phenomenon that can result in a UMAP that is segregated by sample instead of cell type. We find that the biologically-grounded measures produced by our CMS scoring prove to be a robust indicator of cell-type fidelity post data integration, while our modified iLISI scoring is an excellent benchmark of UMAP integration. We demonstrate the ability of this combined CMS and iLISI approach to measuring the most impactful contributors to batch-effect between samples.

Using CMS and iLISI, we reported the ability of common experimental variables to influence the biological interpretation of clusters. In isolation, replicating the library sequencing did not cause any significant loss in cell-type fidelity, while sequencing depth and sample donor both did. However, the effects of sequencing depth were contained entirely within a single, defined cell type while sample donor effects (which include the entangled effects of microfluidic encapsulation and library preparation) were widespread across all cells surveyed. While we identified this phenomenon on PBMCs here, we would expect similar results in other heterogeneous tissues, both with regards to sequencing depth and the effects of sample donor. Pooling PBMC libraries for sequencing together or sequencing libraries independently did not change the observed batch effect. This suggests that pooling libraries for sequencing together may not supply a significant benefit to experimental design or effectively prevent batch-associated systematic variation. We emphasize the critical role of CMS scores in quantifying cell-type fidelity, which helped us to conclude that sample donor is the major contributing variable to batch-effect in the PBMCs evaluated.

While PBMC populations may subtly differ between healthy adult donors, we should be able to pool PBMCs from different donors for the purposes of constructing a reference sample (i.e., a healthy control sample or a cell atlas). To integrate PBMCs from different donors, we applied three popular and often cited batch-correction algorithms^5^ (Harmony^8^, LIGER^11^, and Seurat V3^10^). Although, as expected, we find that all three methods effectively integrated UMAP embeddings, we show that explicit batch-correction may be unnecessary and, in some cases, even harmful. By selecting a suitable normalization and merging strategy through a CMS-guided approach, we produced cell-type integration results comparable to the best batch-correction algorithms. We endorse a supervised approach (such as our CMS-guided optimization) over selecting a batch-correction method for the following reasons: while batch-correction methods do not directly adjust the gene expression matrix, they do affect the dimensional reduction matrix and therefore cluster assignments, which are used to infer cell-type identity. Most significantly, each batch correction method evaluated here produced subtly different cluster assignments from the same data, which could provoke a dangerous potential to misclassify cells and misinterpret results. Therefore, we caution against the application of batch-effect correction methods as a “fix” for systemic batch-effect and place emphasis on the proper, supervised selection of appropriate normalization methods. The choice of the best approach may be aided by a measure such as CMS, or similar, which directly measures the impact of any computational approach on biologically assigned cell-type classifications. By directly quantifying the level of biological signal loss, we can reveal systemic variation that may otherwise go undetected.

To complete our study, we investigated the most symptoms of such systemic variations on the gene expression matrix. We identified the cells most sensitive to misclassification and highlighted a pattern of gene expression which distinguished them from the similar cell type, but correctly classified cells, of the other samples. The differentially expressed genes between the misclassified and correctly classified cells reveal a broad pattern of expression with a low total expression level. We find that, depending on the normalization/merging strategy chosen, these genes may be excluded from the highly variable gene (HVG) list. Strikingly, we show that simple exclusion of these genes significantly improves batch-effect but is insufficient to eliminate all batch-effect in an improperly normalized/merged sample. This suggests that more sources of variation may be present, which cannot be corrected by gene curation alone. We conclude that removing problematic genes *post-hoc* is insufficient and that optimizing the normalization/merging strategy is the best approach to consistently reduce sample-specific variance and highlight biologically relevant signals.

Together, our experiments describe the variables which contribute most to batch-effect (sample donor, sequencing depth) and those which do not significantly contribute (sequencing replicate, sequencing pool). We apply a combined approach of CMS and iLISI scoring to reveal that batch correction algorithms pose a risk to over-merge diverse cell types if not properly supervised, while data normalization offers a simple strategy for minimizing the influence of batch-specific, low expression gene transcripts to the sample. We recognize that while we identify the best strategy to minimize batch-effect in the PBMC samples tested, the optimal strategy may vary by tissue or even by dataset. Rather than endorse a single approach to remove batch-effect, we offer a novel tool, CMS scoring, that, in combination with iLISI can assess cell mismatching and UMAP integration and aid in choosing the correct computational methods for any dataset.

## Materials and Methods

### Sample preparation and single-cell encapsulation

Peripheral venous blood was collected from healthy volunteer donors through Emory University’s Children’s Clinical and Translational Discovery Core. Peripheral Blood Mononuclear Cells (PBMCs) were isolated using the Direct Human PBMC Isolation Kit (StemCell Technologies) according to the manufacturer’s protocol and washed/resuspended in custom RPMI-1640 deficient in biotin, L-glutamine, phenol red, riboflavin, and sodium bicarbonate (defRPMI), and containing 3% newborn calf serum and Benzonase. Cell number and viability were assessed using ViaStain^TM^ Acridine Orange/Propidium Iodide (AOPI, Nexcelom) on a Cellometer K2 cell counter (Nexcelom), following manufacturer recommendations. A maximum of 1×10^6^ cells/donor were stained by oligo-barcoded antibodies (TotalSeq-A or TotalSeq-C; Biolegend, Expedeon) for 30 min on ice, followed by two washes in defRPMI + 0.04% BSA. Cells were resuspended at 1200-1500 cells/ul in defRPMI + 0.04% BSA, passed through a 20uM filter, and counted prior to loading onto a Chromium Controller (10X Genomics). For generating single-cell RNA-seq libraries, cells were loaded to target encapsulation of six thousand cells.

### Library generation and sequencing

Single-cell RNA-seq gene expression and ADT libraries were generated following the manufacturer’s instructions using Chromium Single Cell 3′ Library & Gel Bead Kit v2, with ADT library generation according to (Nat Methods 14, 865–868 (2017) for PBMC samples 1-3 and Chromium Single Cell 5′ Library & Gel Bead Kit v1 with feature barcoding (10X Genomics) for PBMC samples 4 and 5. Gene expression libraries were sequenced to an average depth of 84,098 reads per cell using the Novaseq 6000 platform (Illumina). ADT libraries were sequenced to a target depth of one hundred reads per antibody, per cell, on the Next-seq platform (Illumina). A separate aliquot of cells for each donor was stained with a fluorescent antibody cocktail (Supplementary Table 3) for 30 minutes on ice, followed by washing with defRPMI/3%NBCS + Benzonase (Sigma). Cells were fixed for 30 minutes at RT with FACS lysis solution (BD Biosciences), followed by a final wash and analysis on a BD LSRII or FACSymphony A5 cytometer. Data were analyzed with FlowJo V10. The gating strategy for excluding debris, doublets, and non-viable cells is shown in Supplementary Fig. 3.

### Gene expression (GEX) data processing

Raw sequence data (FASTQ files) was processed in a Linux environment using CellRanger V3 (10X Genomics, version details in Supplementary Table 1) to generate a digital expression matrix. Specifically, splicing-aware aligner STAR was used for sequence alignment to GRCh38 (Ensembl 93). Viable cell barcodes were found automatically by CellRanger. Digital expression matrices were exported for further analyses. Further data analysis was performed in an R environment (Version 3.6.1, CRAN) using the Seurat toolkit (Version 3.2.2, Satija Lab) following previously published workflows ^22^. Following Seurat standard recommendations, data were first filtered for quality using specific QC criteria (maximum mitochondrial content) to limit the analysis to cells with transcriptomes that were not apoptotic. Cell barcodes with mitochondrial transcripts (>5 standard deviations above the median level of mitochondrial transcripts) were suspected to be dying cells and excluded. On average, 99.3% of putative cells met QC criteria and were included in the analysis (range = 99.2 - 99.4).

From the five human PBMC donors (seven libraries), we included a total of 37,442 cells, with a range of 3,758-6,374 cells per donor and a median of 5,729 cells per sample. Genes were selected for differential expression across the sample using Seurat’s highly variable gene selection tool, “FindVariableFeatures.” A setting was chosen to select the 2,000 most variable features identified per sample. Principle Components Analysis (PCA) was used to reduce the dimensionality of the gene expression matrix. A singular value decomposition (SVD) PCA was performed on the subset of highly variable genes. To identify an appropriate number of PCs, we employed a *z*-scoring method. We run a complete PCA reduction and *z*-score the contribution of each PC to the total variance. PCs with z□>□1 were considered significant and used further in the analysis. The SVD PCA returns the right singular values, representing the embeddings of each cell in PC space, and left singular values, representing the loadings (weights) of each gene in the PC space. Cell embeddings (right singular values) were weighted by the variance of each PC.

### Antibody-derived tag (ADT) data processing

ADT data was co-analyzed with GEX data as raw sequence data (FASTQ files) in a Linux environment using Cell Ranger V3 (10X Genomics, version details in Supplementary Table 1). ADT data was aligned directly to a feature reference file containing the sequence of each barcode mapped to the corresponding antibody. Digital expression matrices for ADT protein expression were exported for further analyses. ADT count matrices were normalized and denoised using the R package “dsb” version 0.1.0^23^.

### Data embedding and cell clustering

For 2-dimensional visualization, Uniform Manifold Approximation and Projection (UMAP) reduction was performed on the PCA matrix (https://github.com/lmcinnes/umap). In parallel, independent of the UMAP coordinates, Louvain–Jaccard clustering was performed on the PC-space. This “bottom-up” clustering method employs a stochastic shared-nearest-neighbor (SNN) approach, in which cells are grouped according to their neighbors in PC space. The nearness of two cells is weighted by the Jaccard index or the degree of sharing between the lists of each cell’s nearest neighbors. The algorithm will build small groups of cells and attempt to iteratively merge them into clusters until the modularity is maximized. We found that a resolution of 0.8 was most proper for building biologically meaningful clusters.

### Cell-type identification and classification

Following PCA (performed on the GEX matrix of highly variable genes), we generated clusters by Louvain-Jaccard clustering with a resolution of 0.8, as implemented by the Seurat function “FindClusters”. Using detection of lineage-specific ADT and GEX markers, we assigned each cluster to a single major lineage (markers and representative data depicted in Supplementary Fig. 2). When indicated in the text, differentially expressed genes were determined using the likelihood-ratio test for single-cell gene expression, as implemented in the “bimod” method of the Seurat function “FindMarkers”^10,20^.

### Cell Misclassification Statistic

To generate the Cell Misclassification Statistic (CMS) score, we first assign cell type labels to clusters in a merged, multi-sample dataset. This can be carried out either by following a gating strategy and assigning cell types to entire clusters which fall within each gate or by transferring cluster labels from a previously gated dataset containing shared cells. The transferred labels can be used to aid cell-type identification but should always be confirmed by marker expression following the overall gating strategy. After cell types have been recorded for the merged samples (target dataset), we match a vector of cell identities from the same sample, processed by a different workflow (reference dataset). We then compare the target cell IDs to a vector of the reference cell type IDs. The CMS score generated is the sum of matching values (total number of cell type ID matches) between the two cell-typing replicates, divided by the total number of cells. We then invert the score by subtracting it from one so that the result is a measure of the fraction of cells that change cell type ID between data analysis workflows. CMS can be directly interpreted as the percentage (expressed as a decimal) of cells which have been misclassified by one of the workflows. Where reported, the CMS is the mean score of all samples present.

### Modified Local Inverse Simpson’s Index (LISI)

Our modified LISI score, the integration LISI or iLISI, is an effective indicator of sample mixing in the UMAP. A LISI score represents, for a given cell, the number of neighboring cells which need to be sampled before the same identity class is sampled (in this case, sample ID). In other words, LISI effectively counts how many identity classes are represented in the local neighborhood of each cell^13^. If we merge three samples, then the maximum LISI score will be close to three, and the minimum close to one. However, the actual maximum score may differ by UMAP. To account for differing numbers of samples and to place LISI on a similar scale as CMS, we first take the median LISI score for a merged set of samples using the r package “LISI”^13^ with sample ID used as the cell label. We subtract the minimum LISI score (one) from the median LISI and divide the result by the maximum LISI (the number of samples). The result is an iLISI score on a scale of zero to one, where a score of one is ideal integration and zero is a perfectly segregated sample. We then inverted these scores by subtracting them from 1, such that the best integrated UMAP will achieve a score approaching 0. This was done to align the values with CMS so that higher scores mean better integration across both measurements. iLISI = 1 – (LISI – 1) / (maximum score – 1). Where reported, the iLISI is the mean score of all samples present.

### Data availability

Gene expression and Antibody Derived Tag matrices have been deposited on the Gene Expression Omnibus (GEO).

### Code availability

Full data analysis workflow and R scripts are made available at github.com/Ghosn-Lab/BBabcock

## Acknowledgments

This study was supported in part by Georgia Clinical and Translational Science Alliance (CTSA) through the National Center for Advancing Translational Sciences of the National Institutes of Health under Award number NIH UL1TR002378; Pediatric Research Alliance, Center for Transplantation, and Immune-Mediated Disorders (Children’s Healthcare of Atlanta); and Lowance Center for Human Immunology. We thank Sachin Kumar (Emory University) for helpful conversations. We thank Emory University’s Children’s Clinical and Translational Discovery Core (CCTDC) for providing peripheral blood samples from healthy donors. Flow cytometry data were collected at the Emory’s Pediatrics/Winship Flow Cytometry Core (access supported in part by Children’s Healthcare of Atlanta). Single-cell libraries were sequenced at the Emory Integrated Genomics Core (EIGC), which is subsidized by the Emory University School of Medicine and is one of the Emory Integrated Core Facilities, and at PerkinElmer Genomics Inc.

## Author Contributions

EEBG and BRB conceived the study. BRB performed all computational analyses and developed the CMS scoring method under the supervision of EEBG. BRB, EEBG and AK wrote, reviewed and edited the manuscript. AK performed all tissue processing and flow cytometry, and generated the scRNA-seq libraries. JY performed all data pre-processing workflows, including GEX and ADT alignment. MLW performed flow cytometry analyses. All authors read and approved the final manuscript.

---

**Supplementary Figure 1.**
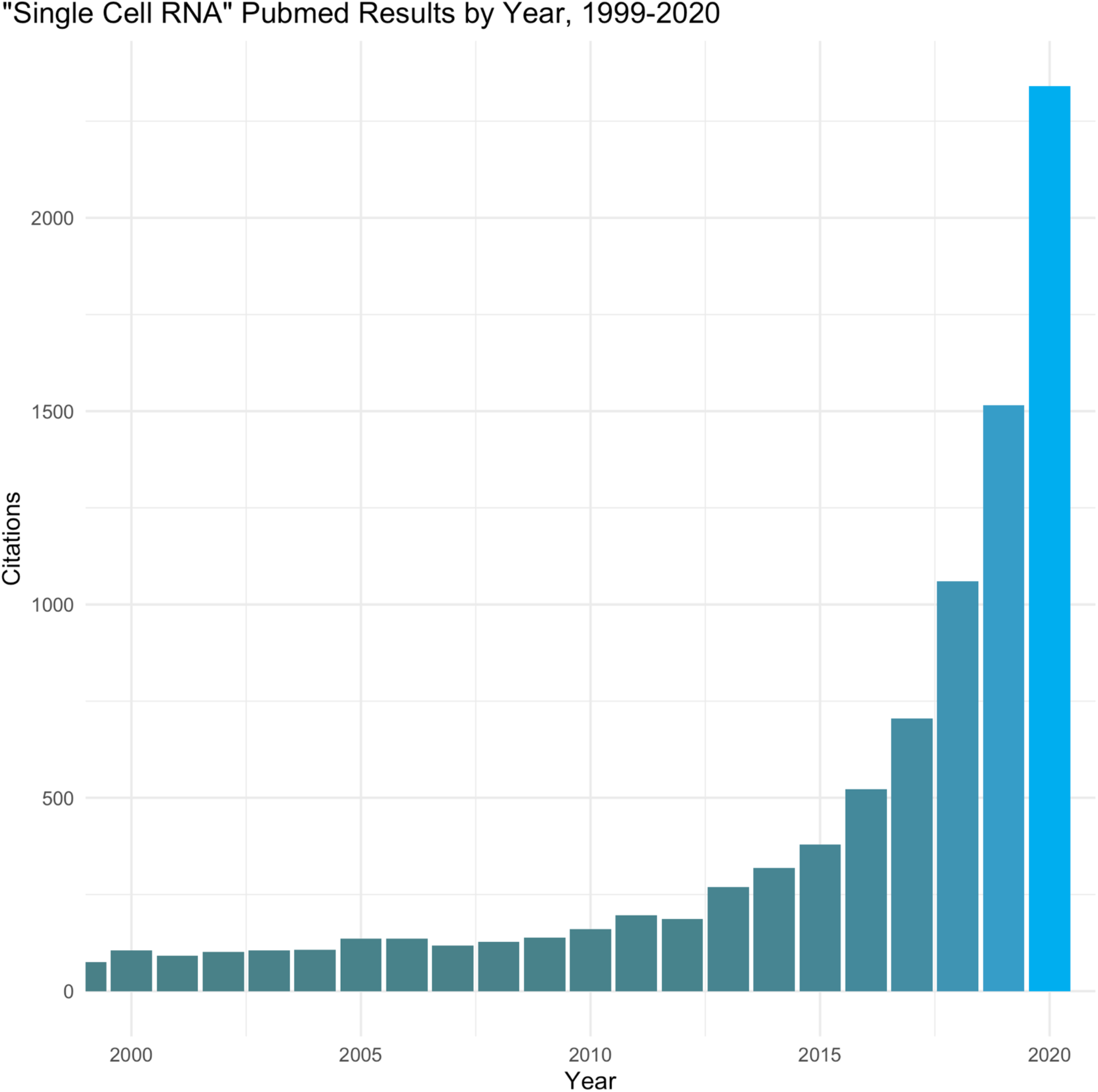
PubMed results for “Single Cell RNA” by year. Increasing trend of publications citing single-cell RNA-seq data in recent decades.

**Supplementary Figure 2.**
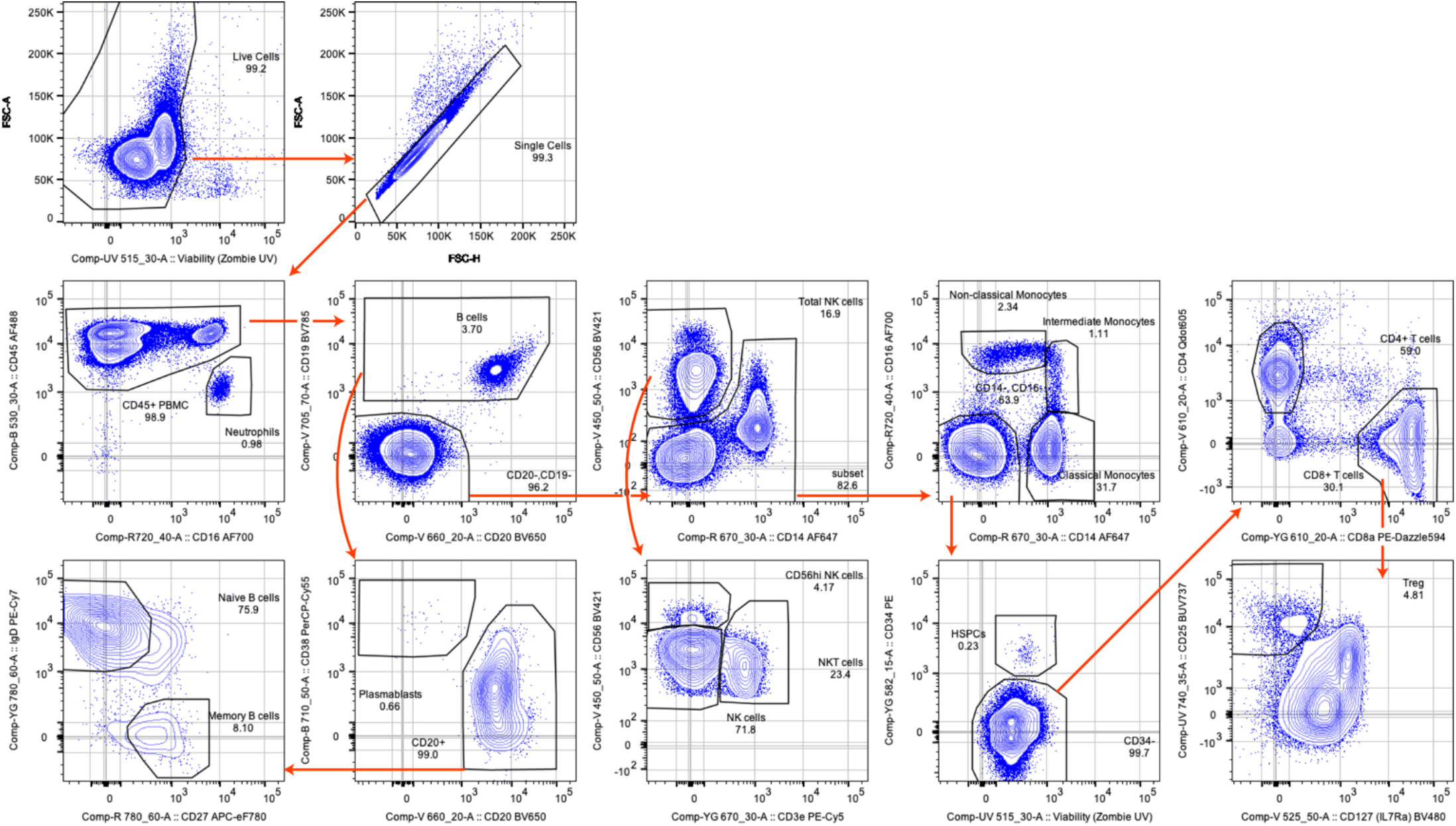
High-dimensional flow cytometry gating strategy to identify major immune lineages in the PBMC samples. Representative gating strategy generated from sample 3, showing markers and gates used to identify major immune cell lineages using an 18-parameter flow cytometry panel.

**Supplementary Figure 3.**
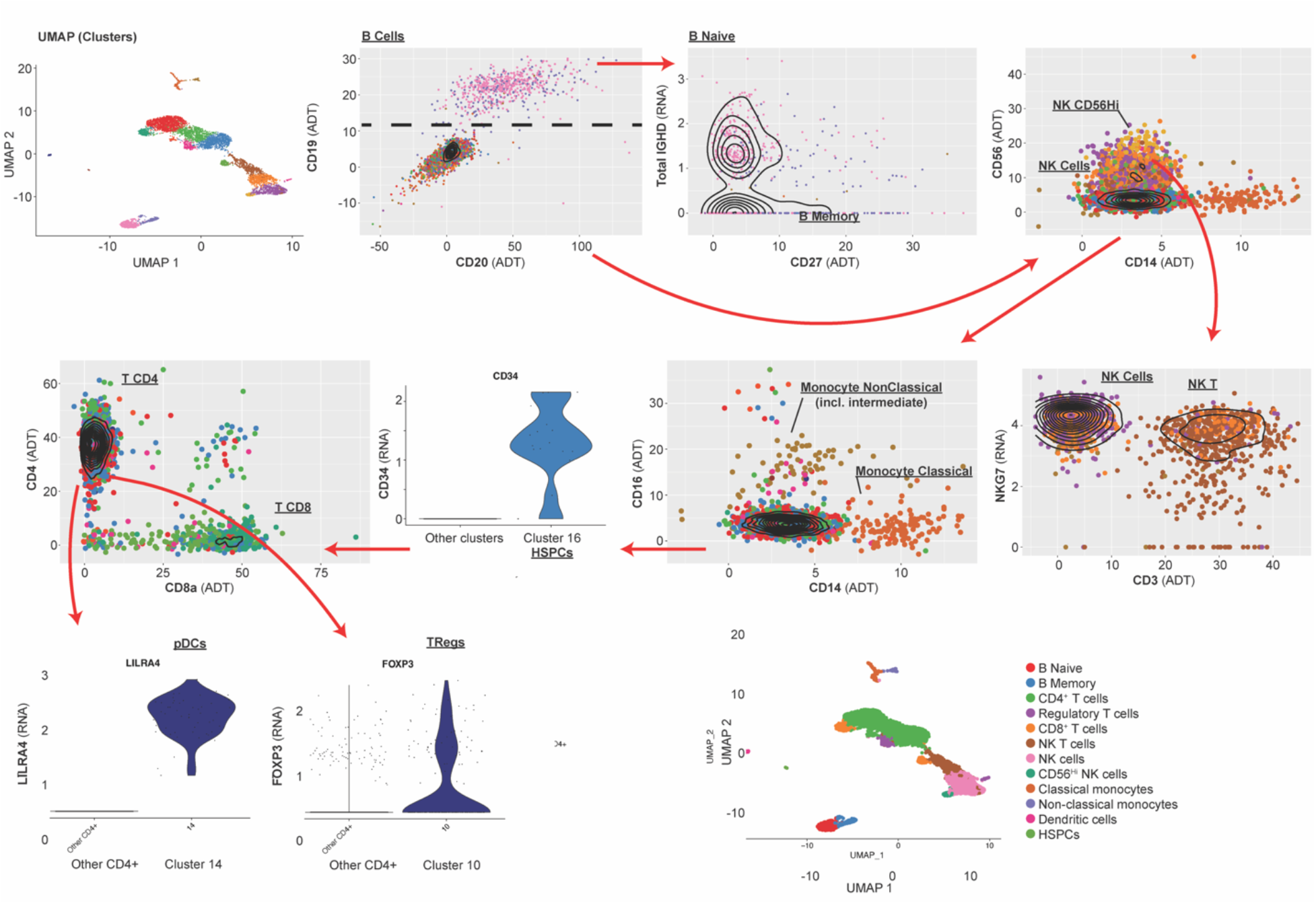
Gating strategy to identify major immune lineages using multi-omics single-cell sequencing data. Representative gating strategy generated from sample 4-A, showing markers and gates used to assign entire clusters to major immune cell lineages based on gene expression (GEX matrix) and cell-surface proteins (ADT matrix).

**Supplementary Figure 4.**
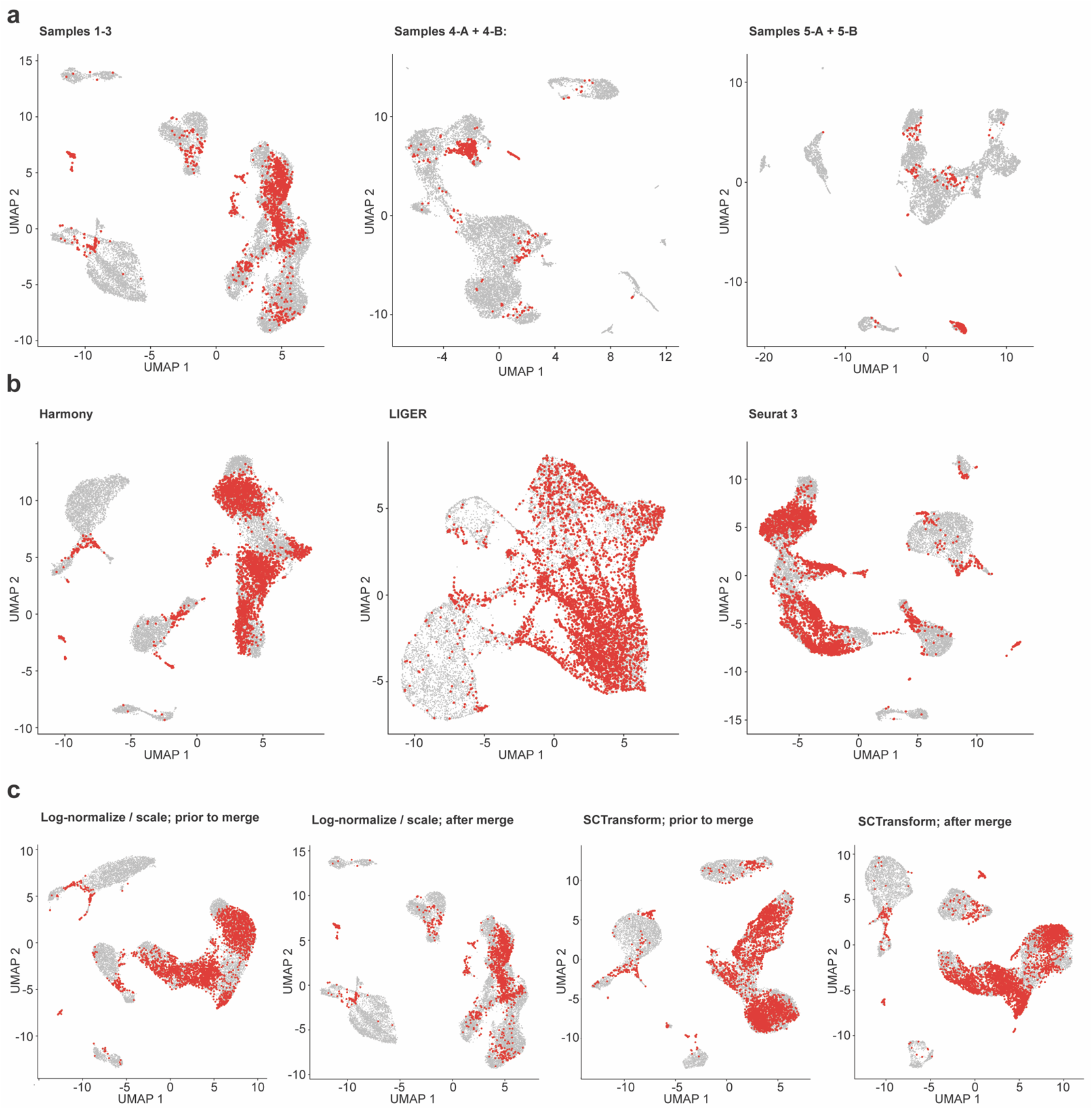
Distribution of misclassified cells after data merging. Red points represent cells that change classification after data merging, as generated by the CMS scoring method. UMAPs and data generated for Figs. 2, 4, and 5.

**Supplementary Figure 5.**
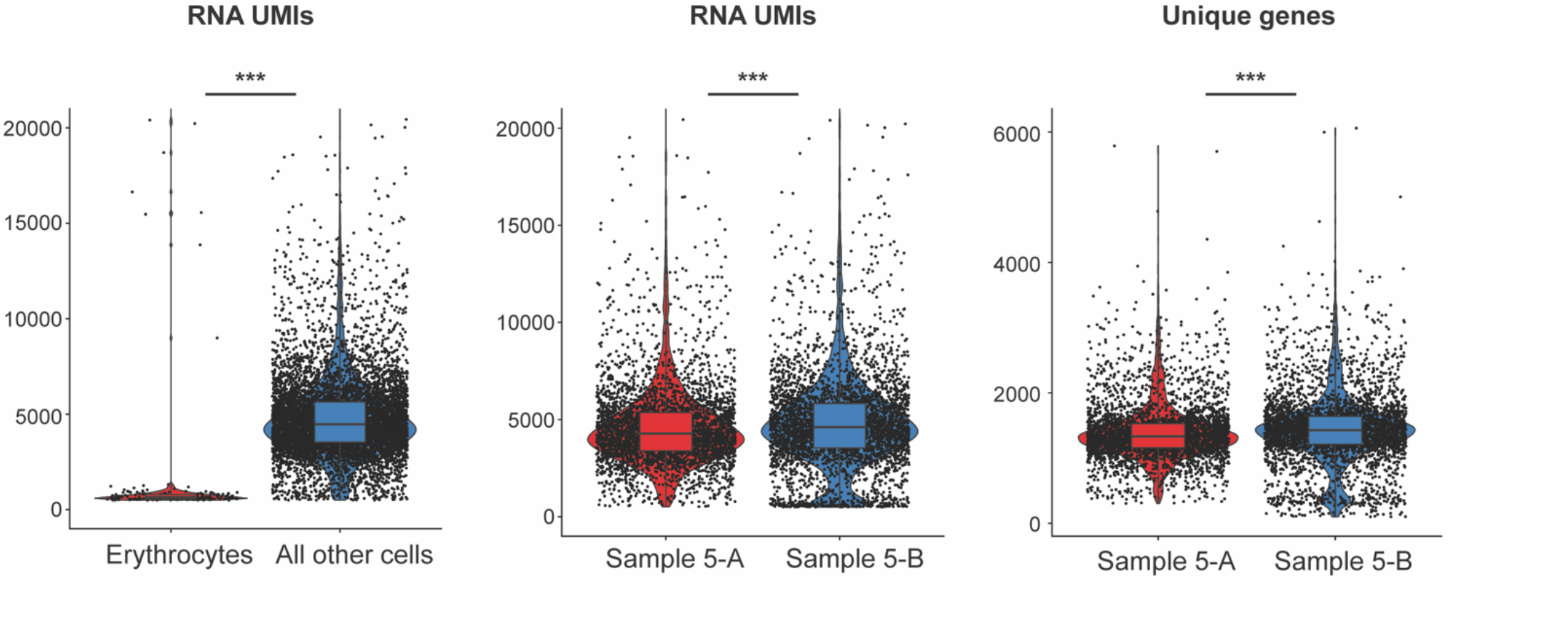
Comparison of sample 5 high- and low-depth sequencing replicates. Comparison of key metrics (number of UMI and number of unique genes) distinguishing high and low read-depth (reads per cell) sequencing replicates of sample 5. Also shown are the number of UMI for sample 5 erythrocytes, as compared to all non-erythrocyte cells. ***p < 1×10^−9^, Wilcoxon rank-sum test.

**Supplementary Table 1.** Donor demographics and library sequencing details.

| Donor ID | Sample ID | Source Tissue | Sex | Age | Isolation method | 10X Chemistry | ADT version | Sequence Platform | Flow cell | CellRanger Version | Reference genome |
| --- | --- | --- | --- | --- | --- | --- | --- | --- | --- | --- | --- |
| Donor 1 | PBMC Sample 1 | PBMC | M | 37 | PBMC isolation SCT kit | 3' v2 | TotalSeq-A | Novaseq | S4-300 | 3.0.1 | GRCh38 (Ensembl 93) |
| Donor 2 | PBMC Sample 2 | PBMC | F | 41 | PBMC isolation SCT kit | 3' v2 | TotalSeq-A | Novaseq | S4-300 | 3.0.1 | GRCh38 (Ensembl 93) |
| Donor 3 | PBMC Sample 3 | PBMC | F | 43 | PBMC isolation SCT kit | 3' v2 | TotalSeq-A | Novaseq | S4-300 | 3.0.1 | GRCh38 (Ensembl 93) |
| Donor 4 | PBMC Sample 4-A | PBMC | M | 46 | PBMC isolation SCT kit | 5' v1 | TotalSeq-C | Novaseq | S4-300 | 3.1.0 | GRCh38 (Ensembl 93) |
| Donor 4 | PBMC Sample 4-B | PBMC | M | 46 | PBMC isolation SCT kit | 5' v1 | TotalSeq-C | Novaseq | S4-300 | 3.1.0 | GRCh38 (Ensembl 93) |
| Donor 5 | PBMC Sample 5-A | PBMC | F | 36 | PBMC isolation SCT kit | 5' v1 | TotalSeq-C | Novaseq | S4-300 | 3.1.0 | GRCh38 (Ensembl 93) |
| Donor 5 | PBMC Sample 5-B | PBMC | F | 36 | PBMC isolation SCT kit | 5' v1 | TotalSeq-C | Novaseq | S4-300 | 3.1.0 | GRCh38 (Ensembl 93) |

| Reads | Cells |
| --- | --- |
| 582,860,987 | 5778 |
| 582,860,987 | 5178 |
| 582,860,987 | 6121 |
| 582,860,987 | 6392 |
| 582,860,987 | 6423 |
| 582,860,987 | 3783 |
| 582,860,987 | 4034 |

**Supplementary Table 2.**
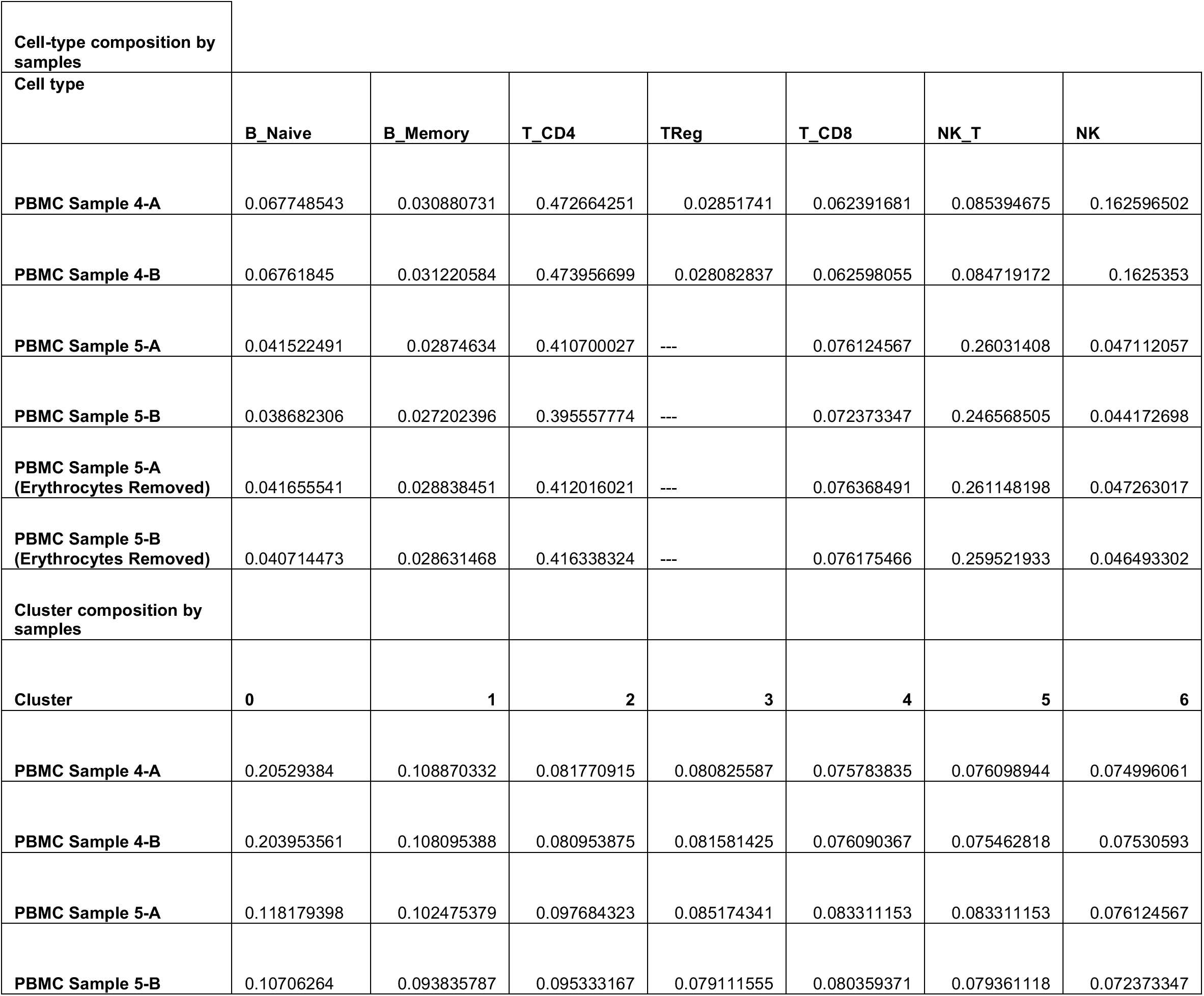
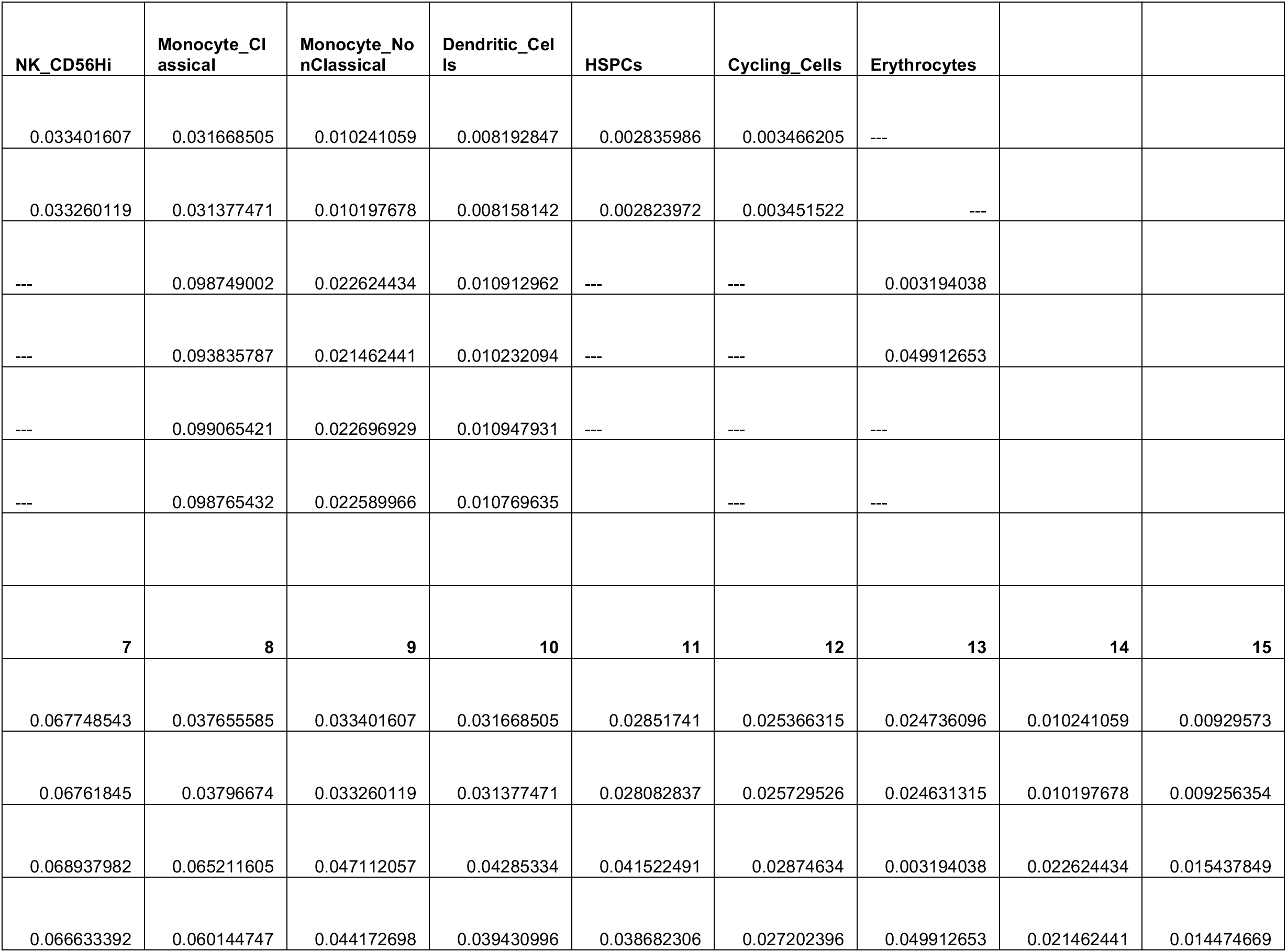
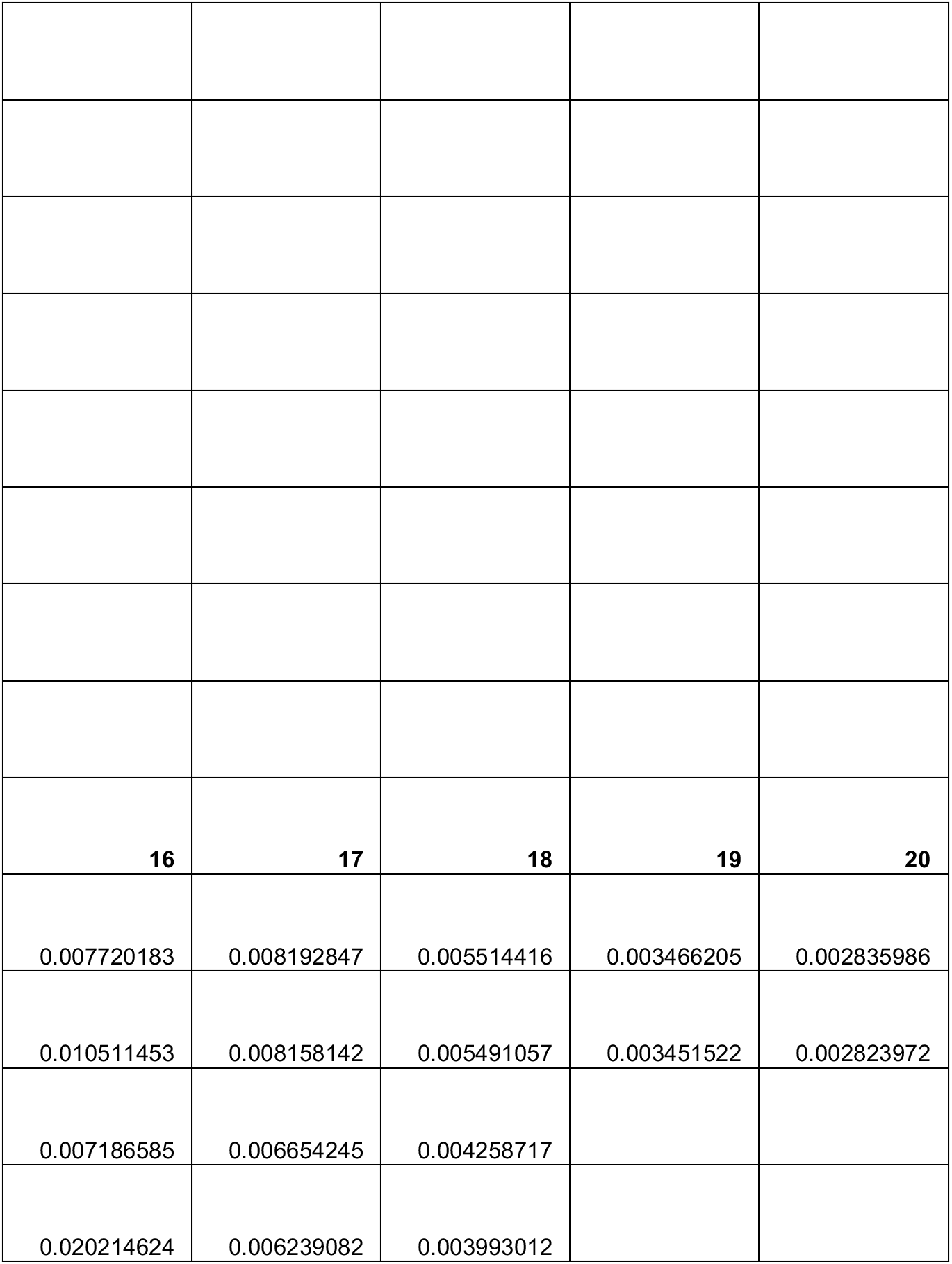
Cell-type and cluster composition (fraction of total sample)

**Supplementary Table 3.**
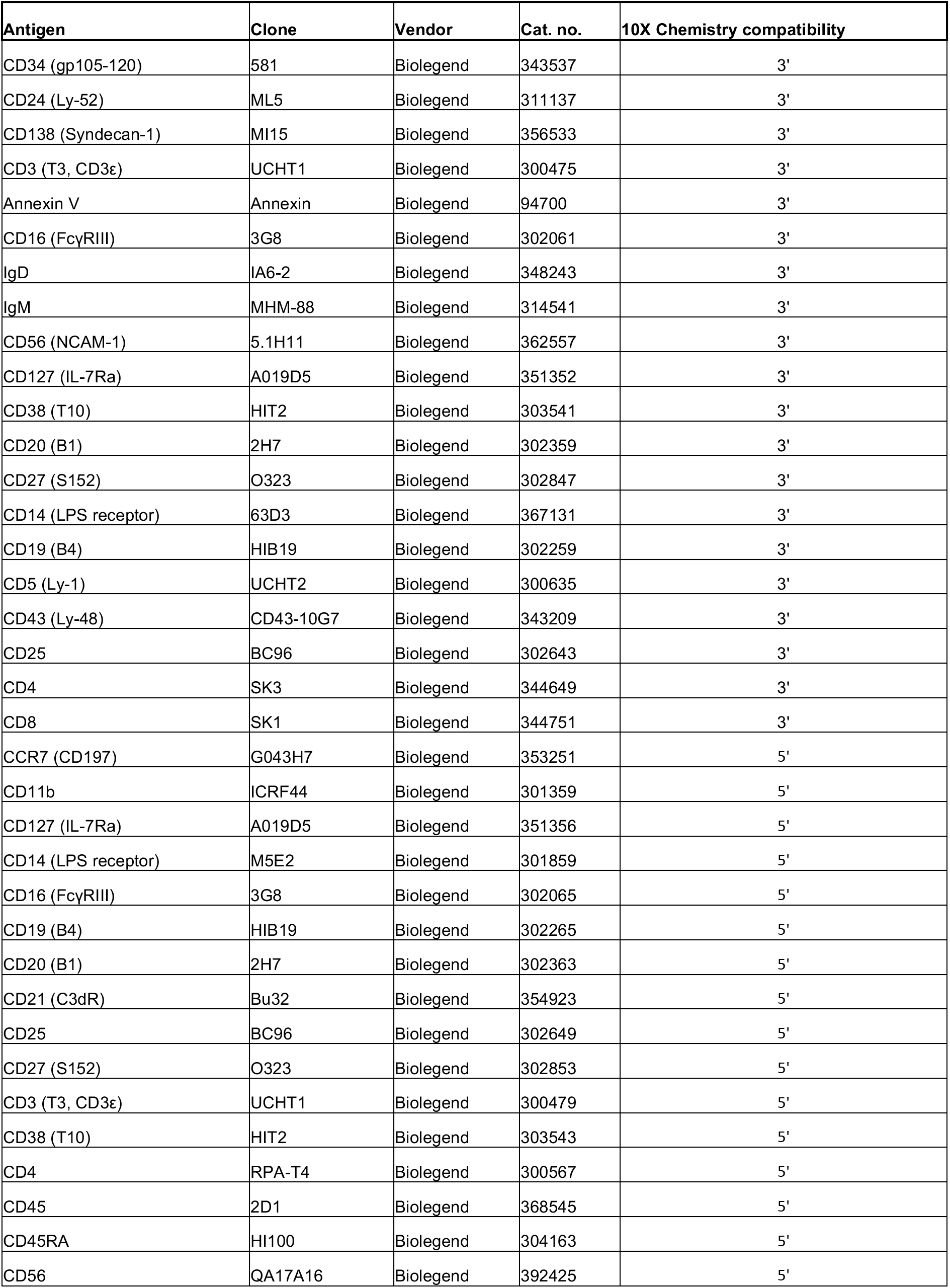
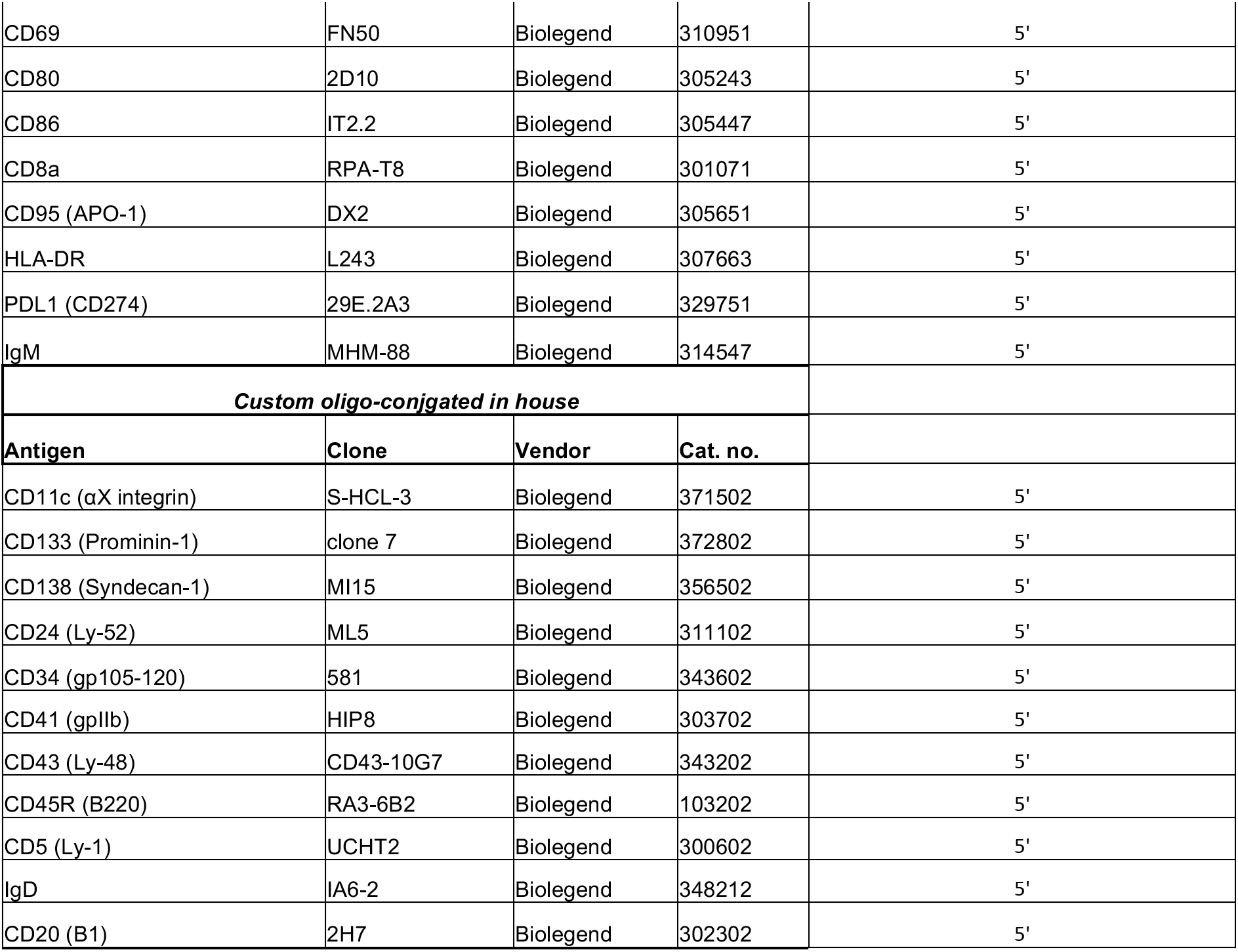
Key reagents and resources.

| Antigen | Clone | Vendor | Cat. no. | 10X Chemistry compatibility |
| --- | --- | --- | --- | --- |
| CD34 (gp105-120) | 581 | Biolegend | 343537 | 3' |
| CD24 (Ly-52) | ML5 | Biolegend | 311137 | 3' |
| CD138 (Syndecan-1) | MI15 | Biolegend | 356533 | 3' |
| CD3 (T3, CD3ε) | UCHT1 | Biolegend | 300475 | 3' |
| Annexin V | Annexin | Biolegend | 94700 | 3' |
| CD16 (FcγRIII) | 3G8 | Biolegend | 302061 | 3' |
| IgD | IA6-2 | Biolegend | 348243 | 3' |
| IgM | MHM-88 | Biolegend | 314541 | 3' |
| CD56 (NCAM-1) | 5.1H11 | Biolegend | 362557 | 3' |
| CD127 (IL-7Ra) | A019D5 | Biolegend | 351352 | 3' |
| CD38 (T10) | HIT2 | Biolegend | 303541 | 3' |
| CD20 (B1) | 2H7 | Biolegend | 302359 | 3' |
| CD27 (S152) | O323 | Biolegend | 302847 | 3' |
| CD14 (LPS receptor) | 63D3 | Biolegend | 367131 | 3' |
| CD19 (B4) | HIB19 | Biolegend | 302259 | 3' |
| CD5 (Ly-1) | UCHT2 | Biolegend | 300635 | 3' |
| CD43 (Ly-48) | CD43-10G7 | Biolegend | 343209 | 3' |
| CD25 | BC96 | Biolegend | 302643 | 3' |
| CD4 | SK3 | Biolegend | 344649 | 3' |
| CD8 | SK1 | Biolegend | 344751 | 3' |
| CCR7 (CD197) | G043H7 | Biolegend | 353251 | 5' |
| CD11b | ICRF44 | Biolegend | 301359 | 5' |
| CD127 (IL-7Ra) | A019D5 | Biolegend | 351356 | 5' |
| CD14 (LPS receptor) | M5E2 | Biolegend | 301859 | 5' |
| CD16 (FcγRIII) | 3G8 | Biolegend | 302065 | 5' |
| CD19 (B4) | HIB19 | Biolegend | 302265 | 5' |
| CD20 (B1) | 2H7 | Biolegend | 302363 | 5' |
| CD21 (C3dR) | Bu32 | Biolegend | 354923 | 5' |
| CD25 | BC96 | Biolegend | 302649 | 5' |
| CD27 (S152) | O323 | Biolegend | 302853 | 5' |
| CD3 (T3, CD3ε) | UCHT1 | Biolegend | 300479 | 5' |
| CD38 (T10) | HIT2 | Biolegend | 303543 | 5' |
| CD4 | RPA-T4 | Biolegend | 300567 | 5' |
| CD45 | 2D1 | Biolegend | 368545 | 5' |
| CD45RA | HI100 | Biolegend | 304163 | 5' |
| CD56 | QA17A16 | Biolegend | 392425 | 5' |
| CD69 | FN50 | Biolegend | 310951 | 5' |
| CD80 | 2D10 | Biolegend | 305243 | 5' |
| CD86 | IT2.2 | Biolegend | 305447 | 5' |
| CD8a | RPA-T8 | Biolegend | 301071 | 5' |
| CD95 (APO-1) | DX2 | Biolegend | 305651 | 5' |
| HLA-DR | L243 | Biolegend | 307663 | 5' |
| PDL1 (CD274) | 29E.2A3 | Biolegend | 329751 | 5' |
| IgM | MHM-88 | Biolegend | 314547 | 5' |
| **Custom oligo-conjugated in house** |  |  |  |  |
| **Antigen** | **Clone** | **Vendor** | **Cat. no.** |  |
| CD11c (αX integrin) | S-HCL-3 | Biolegend | 371502 | 5' |
| CD133 (Prominin-1) | clone 7 | Biolegend | 372802 | 5' |
| CD138 (Syndecan-1) | MI15 | Biolegend | 356502 | 5' |
| CD24 (Ly-52) | ML5 | Biolegend | 311102 | 5' |
| CD34 (gp105-120) | 581 | Biolegend | 343602 | 5' |
| CD41 (gpIIb) | HIP8 | Biolegend | 303702 | 5' |
| CD43 (Ly-48) | CD43-10G7 | Biolegend | 343202 | 5' |
| CD45R (B220) | RA3-6B2 | Biolegend | 103202 | 5' |
| CD5 (Ly-1) | UCHT2 | Biolegend | 300602 | 5' |
| IgD | IA6-2 | Biolegend | 348212 | 5' |
| CD20 (B1) | 2H7 | Biolegend | 302302 | 5' |

## References

1 Yu, P. & Lin, W. Single-cell Transcriptome Study as Big Data. Genomics Proteomics Bioinformatics 14, 21–30, doi:10.1016/j.gpb.2016.01.005 (2016).

2 Luecken, M. D. & Theis, F. J. Current best practices in single-cell RNA-seq analysis: a tutorial. Mol Syst Biol 15, e8746, doi:10.15252/msb.20188746 (2019).

3 Aliverti, E. et al. Projected t-SNE for batch correction. Bioinformatics 36, 3522–3527, doi:10.1093/bioinformatics/btaa189 (2020).

4 Chen, W. et al. A multicenter study benchmarking single-cell RNA sequencing technologies using reference samples. Nat Biotechnol, doi:10.1038/s41587-020-00748-9 (2020).

5 Tran, H. T. N. et al. A benchmark of batch-effect correction methods for single-cell RNA sequencing data. Genome Biol 21, 12, doi:10.1186/s13059-019-1850-9 (2020).

6 Hafemeister, C. & Satija, R. Normalization and variance stabilization of single-cell RNA-seq data using regularized negative binomial regression. Genome Biol 20, 296, doi:10.1186/s13059-019-1874-1 (2019).

7 Hie, B., Bryson, B. & Berger, B. Efficient integration of heterogeneous single-cell transcriptomes using Scanorama. Nat Biotechnol 37, 685–691, doi:10.1038/s41587-019-0113-3 (2019).

8 Korsunsky, I. et al. Fast, sensitive and accurate integration of single-cell data with Harmony. Nat Methods 16, 1289–1296, doi:10.1038/s41592-019-0619-0 (2019).

9 Stein-O’Brien, G. L. et al. Decomposing Cell Identity for Transfer Learning across Cellular Measurements, Platforms, Tissues, and Species. Cell Syst 8, 395–411 e398, doi:10.1016/j.cels.2019.04.004 (2019).

10 Stuart, T. et al. Comprehensive Integration of Single-Cell Data. Cell 177, 1888–1902 e1821, doi:10.1016/j.cell.2019.05.031 (2019).

11 Welch, J. D. et al. Single-Cell Multi-omic Integration Compares and Contrasts Features of Brain Cell Identity. Cell 177, 1873–1887 e1817, doi:10.1016/j.cell.2019.05.006 (2019).

12 Xu, C., et al. Comprehensive multi-omics single-cell data integration reveals greater heterogeneity in the human immune system. bioRxiv (2021).

13 Korsunsky, I. LISI, <https://github.com/immunogenomics/LISI> (2019).

14 Korsunsky, I. How to use Harmony with Seurat V3, <https://github.com/immunogenomics/harmony/blob/master/docs/SeuratV3> (2019).

15 Welch, J. D. LIGER, <https://github.com/welch-lab/liger> (2021).

16 Butler, A. Integrating Seurat objects using LIGER, <https://github.com/satijalab/seurat-wrappers/blob/master/docs/liger.md> (2021).

17 Satija, R. Integration and Label Transfer: SCTransform Vignette, <https://satijalab.org/seurat/archive/v3.0/integration.html> (2019).

18 Lytal, N., Ran, D. & An, L. Normalization Methods on Single-Cell RNA-seq Data: An Empirical Survey. Front Genet 11, 41, doi:10.3389/fgene.2020.00041 (2020).

19 Hou, W., Ji, Z., Ji, H. & Hicks, S. C. A systematic evaluation of single-cell RNA-sequencing imputation methods. Genome Biol 21, 218, doi:10.1186/s13059-020-02132-x (2020).

20 McDavid, A. et al. Data exploration, quality control and testing in single-cell qPCR-based gene expression experiments. Bioinformatics 29, 461–467, doi:10.1093/bioinformatics/bts714 (2013).

21 Rousseeuw, P. J. Silhouettes: A graphical aid to the interpretation and validation of cluster analysis. Journal of Computational and Applied Mathematics 20, 53–65, doi:10.1016/0377-0427(87)90125-7 (1987).

22 Ocasio, J. et al. scRNA-seq in medulloblastoma shows cellular heterogeneity and lineage expansion support resistance to SHH inhibitor therapy. Nat Commun 10, 5829, doi:10.1038/s41467-019-13657-6 (2019).

23 Mulè, M. P., Martins, A. J. & Tsang, J. S. Normalizing and denoising protein expression data from droplet-based single cell profiling. bioRxiv, doi:https://doi.org/10.1101/2020.02.24.963603 (2021).

